# Alternative splicing liberates a cryptic cytoplasmic isoform of mitochondrial MECR that antagonizes influenza virus

**DOI:** 10.1101/2020.11.09.355982

**Authors:** Steven F. Baker, Helene Meistermann, Manuel Tzouros, Aaron Baker, Sabrina Golling, Juliane Siebourg Polster, Mitchell P. Ledwith, Anthony Gitter, Angelique Augustin, Hassan Javanbakht, Andrew Mehle

## Abstract

Viruses must balance their reliance on host cell machinery for replication while avoiding host defense. Influenza A viruses are zoonotic agents that frequently switch hosts, causing localized outbreaks with the potential for larger pandemics. The host range of influenza virus is limited by the need for successful interactions between the virus and cellular partners. Here we used immuno-competitive capture-mass spectrometry to identify cellular proteins that interact with human- and avian-style viral polymerases. We focused on the pro-viral activity of heterogenous nuclear ribonuclear protein U-like 1 (hnRNP UL1) and the anti-viral activity of mitochondrial enoyl CoA-reductase (MECR). MECR is localized to mitochondria where it functions in mitochondrial fatty acid synthesis (mtFAS). While a small fraction of the polymerase subunit PB2 localizes to the mitochondria, PB2 did not interact with full-length MECR. By contrast, a minor splice variant produces cytoplasmic MECR (cMECR) that interacts with PB2. cMECR binds the viral polymerase and suppresses viral replication by blocking assembly of viral ribonucleoprotein complexes (RNPs). MECR ablation through genome editing or drug treatment is detrimental for cell health, creating a generic block to virus replication. Using the yeast homolog Etr1 to supply the metabolic functions of MECR in MECR-null cells, we showed that specific antiviral activity is independent of mtFAS and lies solely within cMECR. Thus, alternative splicing produces a cryptic antiviral protein that is embedded within a key metabolic enzyme.

## Introduction

To move from one host to the next, viruses must overcome cross-species transmission barriers by engaging divergent cellular co-factors while evading host-encoded antiviral proteins. Emerging and re-emerging influenza viruses regularly surmount host barriers through adaptive mutations that allow them to interface with new host cell environments. Migratory waterfowl are the natural host reservoir for influenza A viruses. Spillover into new hosts and adaptation has led to endemic infection in mammals including humans, pigs, dogs, and horses. Yearly influenza virus epidemics shape public health programs worldwide. Protection through vaccination and treatment with antivirals helps slow influenza virus, yet infections still cause ∼61,000 deaths yearly in the United States during high severity seasons^1^. Thus, it is paramount to understand what conserved cellular functions are engaged by viruses and allow them to establish infection, especially during initial cross-species infections.

Viruses are completely dependent upon cellular co-factors. The host counters this dependence and maintains pressure on the invading virus through positive selection of mutations on critical co-factors or antiviral proteins. The virus responds with its own set of adaptive mutations. The recursive process of host evasion countered by viral adaptation establishes a so-called molecular arms race, or Red Queen genetic conflict^2^. The process is aided by the fact that most host antiviral genes are non-essential for the host cell, enabling mutation without compromising viability, and mutational tolerance is further bolstered through gene duplication and transcriptional regulation^3^. Geneduplication and diversification allows hosts to mutate otherwise essential genes whereas changes in transcriptional regulation can selectively activate genes should their constitutive expression be detrimental. To counter mutational tolerance, it has been suggested that viruses target essential genes as host co-factors, because the genes are less prone to variability^4^.

Due to their rapid replication and high mutation rates relative to the host, viruses are eventually successful in both winning genetic conflicts and adapting to new hosts. For example, the human protein ANP32A or its paralog ANP32B are required for influenza viral genome replication^5^. While ANP32A/B double knockout mice are not viable, functional overlap between both proteins theoretically allows hosts to test virus escape mutations in one or the other paralog without a strong global fitness cost^6^. This is perhaps exemplified by the insertion of duplicated sequence in the avian *ANP32A* locus and a loss-of-function mutation in *ANP32B*. This has forced avian-adapted influenza polymerases to adapt and become solely reliant upon ANP32A in most avian hosts^7,8^. As a consequence, avian-adapted viral polymerases function poorly in mammals^9^. Nonetheless, a single amino acid change in the viral polymerase PB2 subunit (E627K) allows avian influenza viruses to rapidly adapt to and exploit human ANP32A^10^.

The viral ribonucleoprotein complex (RNP) is the minimal unit for viral genome replication and a major hotspot for influenza virus host-adaptation. RNPs are helically wound structures composed of genomic RNA encapsidated by viral nucleoprotein (NP) with the heterotrimeric RNA-dependent RNA polymerase at one end binding both the 5’ and 3’ termini of the genome. The polymerase is composed of the PB1, PB2 and PA subunits. All of the enzymatic activities required for replication and transcription are intrinsic to the polymerase, whereas host factors serve as essential co-factors or modulate RNP function^11,12^. The polymerase assumes distinct conformations during each step of the replication or transcription cycle, presenting unique interfaces for host protein interactions (reviewed in ^13,14^). Indeed, MCM, ANP32A or ANP32B, and RNAP2 are host proteins that facilitate replication of the plus-sense genome intermediate cRNA, the minus-sense genomic vRNA, or transcription, respectively^5,15,16^.

Understanding which host proteins the polymerase requires for replication, especially those that it engages when influenza virus jumps from one host to the next, is essential for understanding the genetic conflicts that establish barriers to cross-species transmission. Transmission of influenza virus from birds to humans requires that the incoming vRNPs successfully interact with cellular factors to, at a minimum, provide sufficient levels of replication during which adaptive mutations can arise. To understand how avian viruses engage the foreign intracellular environment of human cells, we used immuno-competitive capture-mass spectrometry (ICC-MS) to define interaction networks between human proteins and avian or human-adapted viral PB2. We identified heterogenous nuclear ribonuclear protein U-like 1 (hnRNP UL1) as a cellular co-factor that supports influenza virus replication and mitochondrial enoyl CoA-reductase (MECR) that plays an anti-viral role. MECR localizes to the mitochondria and is a critical enzyme in mitochondrial fatty acid synthesis (mtFAS), raising questions as to how it could counteract the viral polymerase in the cell nucleus. Surprisingly, we demonstrated that MECR antiviral activity is derived from a cryptic splice isoform that produces cytosolic MECR (cMECR). cMECR is identical to MECR other than its lack of a mitochondrial targeting sequence. cMECR interferes with viral infection by inhibiting *de novo* assembly of vRNPs. Loss of MECR cripples mtFAS and cellular metabolism, resulting in defects in cell health and a generic block to viral replication. However, repairing MECR deficient cells with the yeast homolog Etr1, which lacks a cMECR-like isoform, showed that cMECR alone suppresses influenza virus replication independent of mtFAS. Thus, *MECR* encodes a critical metabolic enzyme important for cell health while also concealing the antiviral protein cMECR that is revealed by differential splicing.

## Results

### Identification of host proteins and pathways that interface with influenza virus polymerase

During zoonoses, avian influenza virus polymerases must co-opt mammalian host processes and proteins to direct virus replication. To identify polymerase co-factors important for zoonotic and endemic transmission, we infected human lung cells with virus encoding avian-style PB2 E627 or the human style PB2 K627 and performed affinity purification-MS (AP-MS) for PB2. As these experiments were performed using infected cells, PB2 exists alone, as part of the heterotrimeric polymerase, or as part of the larger RNP. AP-MS approaches can be complicated by high levels of nonspecific interactions. We overcame these limitations by performing ICC-MS (**Fig. 1a**). ICC-MS is a label-free strategy that includes a competition step with in-solution antibody prior to affinity purification using antibody already bound to a solid support. ICC-MS thus distinguishes specific interactions, which can be competed, from nonspecific interactions, which are unaffected by the competition step^17^. The competition profiles are further used to rank-order putative host interactors. Increasing amounts of competing antibody specifically reduced capture of PB2 and the viral RNP (**Fig. 1b**). Avian-style PB2 E627 was more sensitive to competition, possibly because this virus is restricted in human cells and expresses lower levels of viral proteins^18^. The samples were then processed for protein identification by LC-MS/MS. Effective competition was confirmed where increased competitor antibody decreased the relative abundance of PB2 itself, as well as the RNP components PB1, PA, and NP that interact with PB2 (**Fig. 1c and Supp. Fig. 1a**). 1,377 proteins were identified in precipitates from PB2 K627 or E627 infections at all five antibody concentrations and in at least two of three biological replicates. Hits were prioritized based on their competition curve profiles to produce a focused list of 22 candidates (**Supp. Table 1**). We also included ANP32A, EWSR1, FUS, and GMPS that were identified in a pilot screen, but not among the top candidates in the subsequent analysis. Several of the PB2 ICC-MS interactors, including ADAR, ANP32A, ATP7A, and KPNA3, were previously identified in proteomic screens for influenza virus polymerase co-factors and studied in detail, providing confidence in our approach^5,19–21^. The majority of PB2 interactors were found during infection with avian- or human-style polymerases, suggesting the identification of co-factors conserved for both zoonotic and endemic infections.

**Fig. 1:**
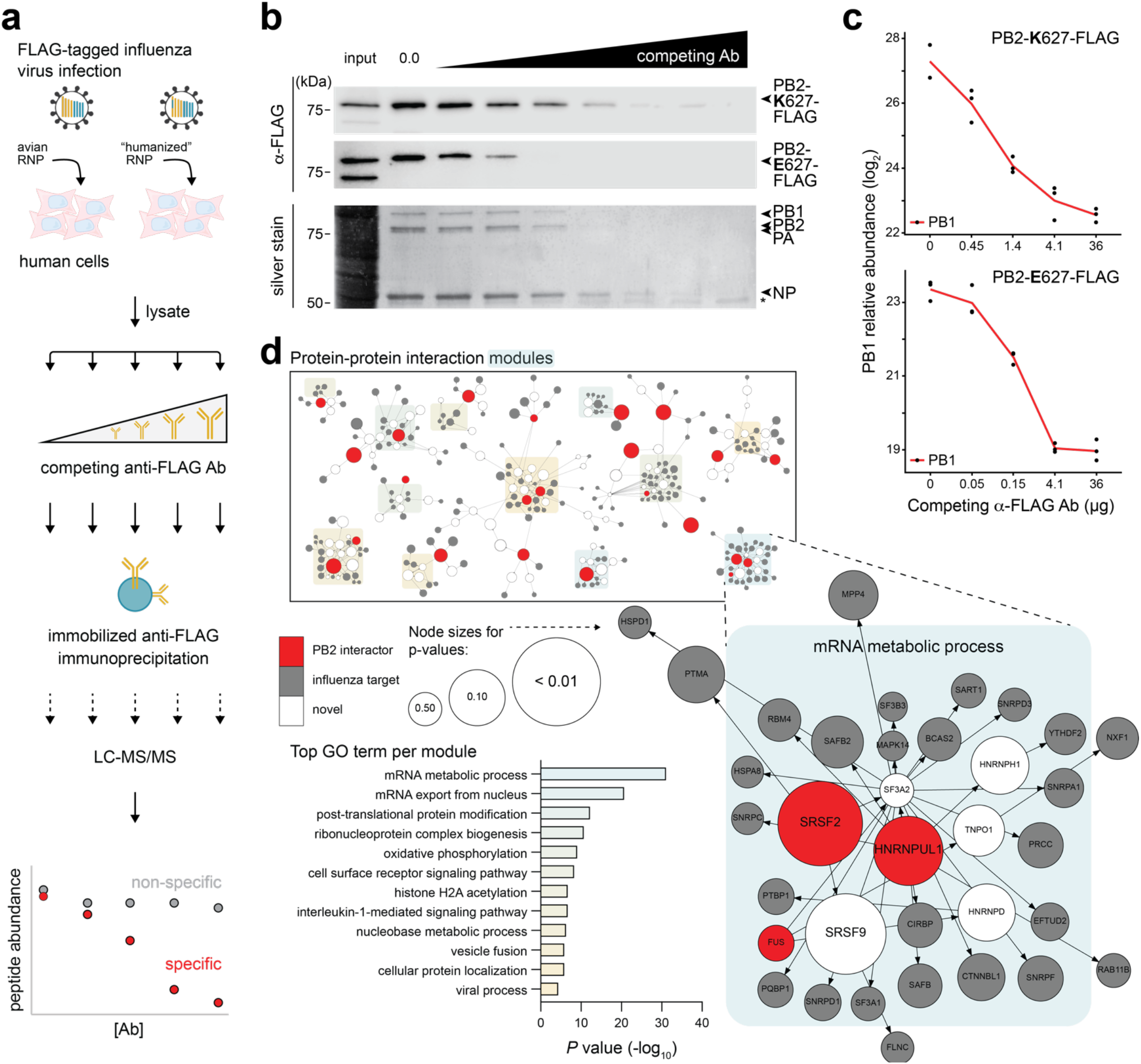
ICC-MS defines an influenza polymerase interactome. **a**, Schematic diagram to identify PB2 interactome. Lysate fractions from human A549 cells infected with PB2-FLAG tagged influenza virus encoding avian RNP or the “humanized” PB2-E627K RNP were incubated with competing soluble anti-FLAG antibody followed by capture with resin-bound anti-FLAG and LC-MS/MS. **b-c**, Immuno-competitive capture of the viral RNP. **b**, Detection of PB2 627E and 627K (*top*: western blot) or PB2 627K with co-precipitating RNP components (*bottom*: silver stain). *, IgG heavy chain. **c**, Relative protein abundance of PB1 in PB2 ICC-MS samples shows decreasing capture with increasing competition antibody. Data shown are in biological triplicate. **d**, Minimum-cost flow simulations connect top PB2 interactors identified by ICC-MS (red) to previously identified influenza host factors (gray), in some cases through other host proteins (white). Modules comprising different PB2 interactors were enriched for GO terms, the most significant of which is enlarged. Node sizes indicate empirical *P* values derived from the control flow simulations.

To increase the power of our ICC-MS results, we performed protein-protein interaction network analyses tailored for influenza virus. Networks were constructed of experimentally demonstrated protein-protein interactions (STRING; ^22^). We then performed minimum-cost flow simulations where the candidate PB2 interactors served as sources to link to targets marked as influenza host factors based on data from six genome-wide screens^23–29^. 23 of the 26 PB2 interactors identified by ICC-MS readily formed subnetworks with previously identified influenza host factors and revealed key cellular processes defined by GO enrichment scores for each of the 12 subnetworks (**Fig. 1d; Supp. Fig. 1b; Supp. Table 2a-b**). Further, they uncovered new protein partners within these subnetworks and connected to highly significant modules not previously associated with influenza virus.

Flow simulations can be biased due to nodes forming spurious links in order to reach large multi-partner targets. But, as detailed in the Methods, the PB2 subnetworks were specific to simulations programmed with PB2 interactors and based on influenza virus host factors (**Supp. Table 2**). To further test the networks, we queried them using host proteins with thoroughly studied interactions as sources: PKR and RIG-I, proteins important for innate immune defense against influenza virus (reviewed in ^30,31^); CRM1 and NXF1, proteins important for viral nuclear-cytoplasmic transport^32,33^; and, EXOSC3 and UBR4, proteins that were identified through rigorous omics approaches^4,34^. These six proteins were used as sources to connect to influenza host factor targets, then controlled through two separate simulations as above. Both flow simulations produced similar results that recapitulate *in silico* the biochemically defined interaction networks, and suggest new proteins that may be important for these processes (**Supp. Fig. 1d**; **Supp. Table 2c**,**d**).

### *HNRNPUL1* promotes and *MECR* restricts viral infection

To test the functional role of proteins identified by ICC-MS, candidate interactors were knocked down by siRNA treatment in A549 cells prior to infection with influenza virus (**Fig. 2a**). Infections were performed with human-style PB2 K627 and avian-style E627 viruses to detect any species-specific dependence. NXF1, an essential host co-factor, was knocked down as a positive control and caused a severe decrease in replication, as expected^24^. Knockdown of hnRNP UL1 reduced viral titers to ∼25% of the non-targeting control, whereas knockdown of MECR significantly increased titers 2.5 to 5-fold. Knockdown of other interactors had only modest effects in A549 cells, possibly because these factors function redundantly (e.g. importin-α isoforms, ANP32A and ANP32B), are needed in very limited quantities (e.g. ANP32A), or are not essential for viral replication^20,35^. Titers in the knockdown cells for virus encoding PB2 K627 or PB2 E627 were highly correlated (Pearson’s *r* ^2^ = 0.791), suggesting our interactors had comparable roles for human-signature or avian-signature virus polymerases (**Fig. 2b**).

**Fig. 2:**
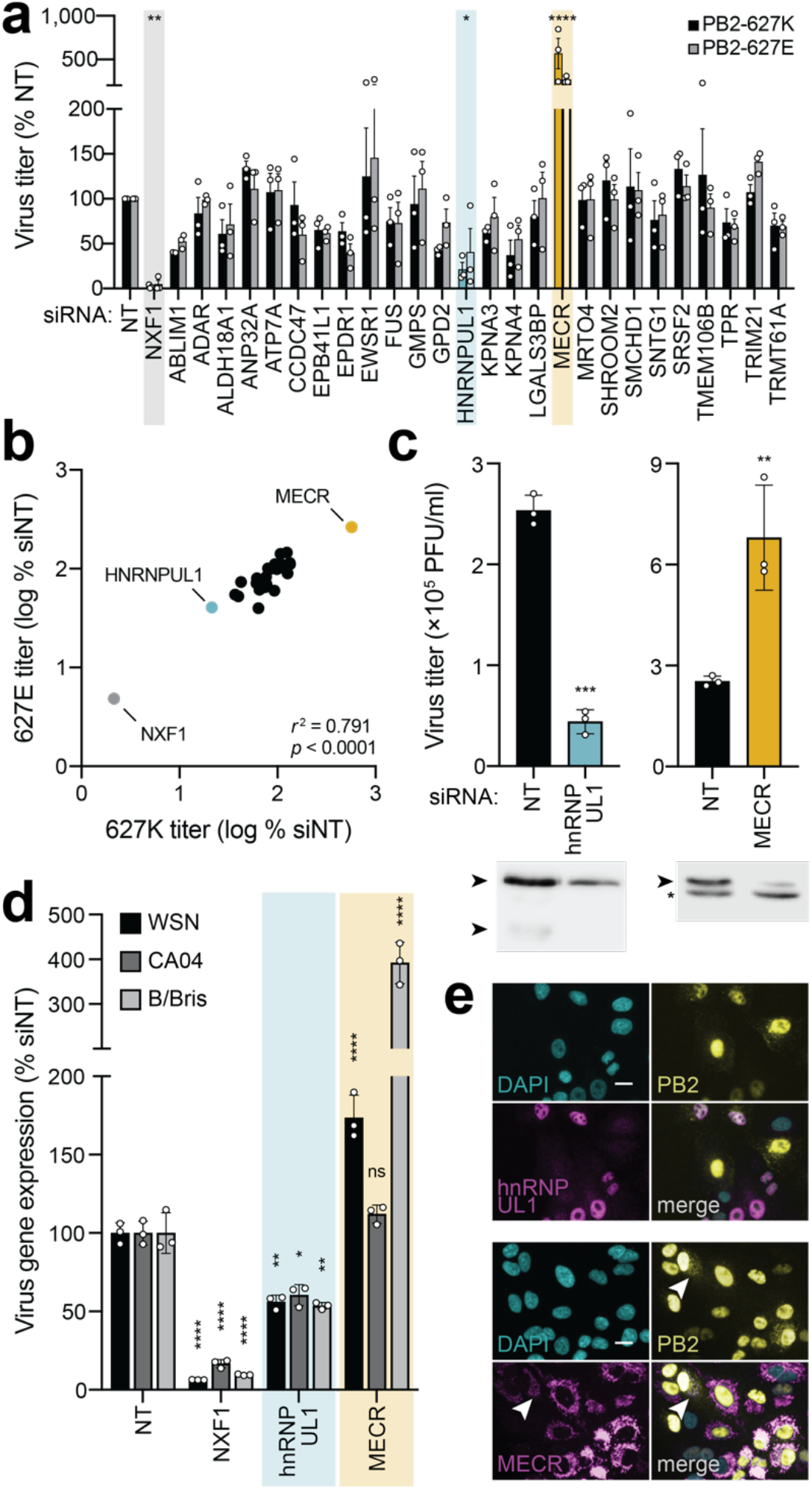
Functional analysis of top candidate PB2 interactors reveals important roles for hnRNP UL1 and MECR. **a**, Secondary screening of proteomic hits by siRNA treatment and reporter virus infection. After knockdown, A549 cells were infected with human (PB2-627K; MOI, 0.01) or avian-adapted (PB2-627E; MOI, 0.05) WSN NLuc virus for 24 h. Viral supernatants were titered and normalized to a non-targeting control (NT). Control NXF1 (gray) and outliers hnRNP UL1 (cyan) and MECR (yellow) are highlighted. **b**, Concordance of virus titer for PB2-627E *vs* PB2-627K virus infections in siRNA-treated cells (from **a**). Statistical analysis performed with a two-tailed Pearson correlation coefficient. **c**, Multicycle virus replication of WT virus was measured 24 h post infection in A549 cells treated with the indicated siRNAs. Knockdown efficiency was analyzed by western blot. Asterisk indicates non-specific band. **d**, Knockdown impacts viral gene expression of divergent influenza viruses. siRNA-treated A549 cells were infected with reporter viruses based on WSN (pre-2009 H1N1; MOI, 0.1), CA04 (pandemic 2009 H1N1; MOI, 0.5), or B/Bris (Victoria-lineage; MOI, 1). Viral gene expression was measured 8 h post infection and normalized to NT controls. **e**, A549 cells stably expressing hnRNP UL1 (*top*) or MECR (*bottom*) were infected with WSN PB2-FLAG (MOI 3, 8 h). Protein localization was detected by immunofluorescence and nuclei were visualized with DAPI. Arrow indicates minor PB2 population consistent with previously reported patterns of mitochondrial localization. Scale bars, 20 µm. Data in **a** are mean ± SEM of *n* = 3 biological replicates. Comparisons were performed with two-way ANOVA with *post hoc* Fisher’s LSD test. For **c** and **d**, data are mean ± SD of *n* = 3. Comparisons were performed with a two-tailed Student’s t test (**c**) or a two-way ANOVA with *post hoc* Dunnett’s multiple comparisons test (**d**); *, *P* < 0.05; **, *P* < 0.01; ***, *P* < 0.001; ****, *P* < 0.0001; ns, not significant.

Separate experiments confirmed knockdown of hnRNP UL1 and MECR (**Fig. 2c**). Reductions in hnRNP UL1 protein levels decreased viral replication, whereas reduction in MECR protein levels increased titers. Similar knockdown phenotypes were detected during multicycle replication in another human cell line, 293T (**Supp. Fig. 2a**,**b**). We extended these findings to primary isolates of influenza A virus from the 2009 pandemic (A/California/04/2009 [CA04]; H1N1) and influenza B virus (B/Brisbane/60/2008 [B/Bris]) (**Fig. 2d**). Loss of hnRNP UL1 reduced viral gene expression during infection for all strains, as did our control target NXF1. Knockdown of MECR again increased infection by WSN, but not CA04. B/Bris was even more impacted by MECR knockdown with a ∼4-fold increase in viral gene expression compared to 1.5 to 2-fold effect seen for WSN. We also measured viral gene expression during a single round of infection. Similar to results with viral replication, viral gene expression was reduced when hnRNP UL1 was knocked down and increased when MECR was knocked down, and this was independent of the identity of PB2 residue 627 (**Supp. Fig. 2c**,**d**). These data suggest that hnRNP UL1 functions as a pro-viral factor, contrasting with the anti-viral activity associated with MECR expression.

### hnRNP UL1 interacts with the viral replication machinery to promote replication

hnRNP UL1 is an RNA-binding protein that plays a role in nucleocytoplasmic RNA transport as well as DNA end resection signaling during double strand break repair^36,37^. As a member of the hnRNP family of proteins that are well-characterized regulators of pre-mRNA processing, hnRNP UL1 has been shown to interact with NXF1 and NS1-BP, proteins that help coordinate export of influenza viral mRNAs and splicing of the viral genome, respectively^33,38,39^. Consistent with its known function, immunofluorescence assays showed that hnRNP UL1 is present primarily in the nucleus of infected cells where it co-localized with PB2 (**Fig. 2e)**. Infection does not appear to change hnRNP UL1 localization. We tested interactions between the viral polymerase and endogenous hnRNP UL1. Cells were infected with WT virus or virus encoding PB2-FLAG and subject to FLAG immunoprecipitation. Endogenous hnRNP UL1 co-precipitated in the presence of PB2-FLAG, as did the polymerase subunit PB1, but neither were present in the control precipitation with untagged PB2 (**Fig. 3a)**, demonstrating specific association with PB2 and confirming our ICC-MS.

**Fig. 3:**
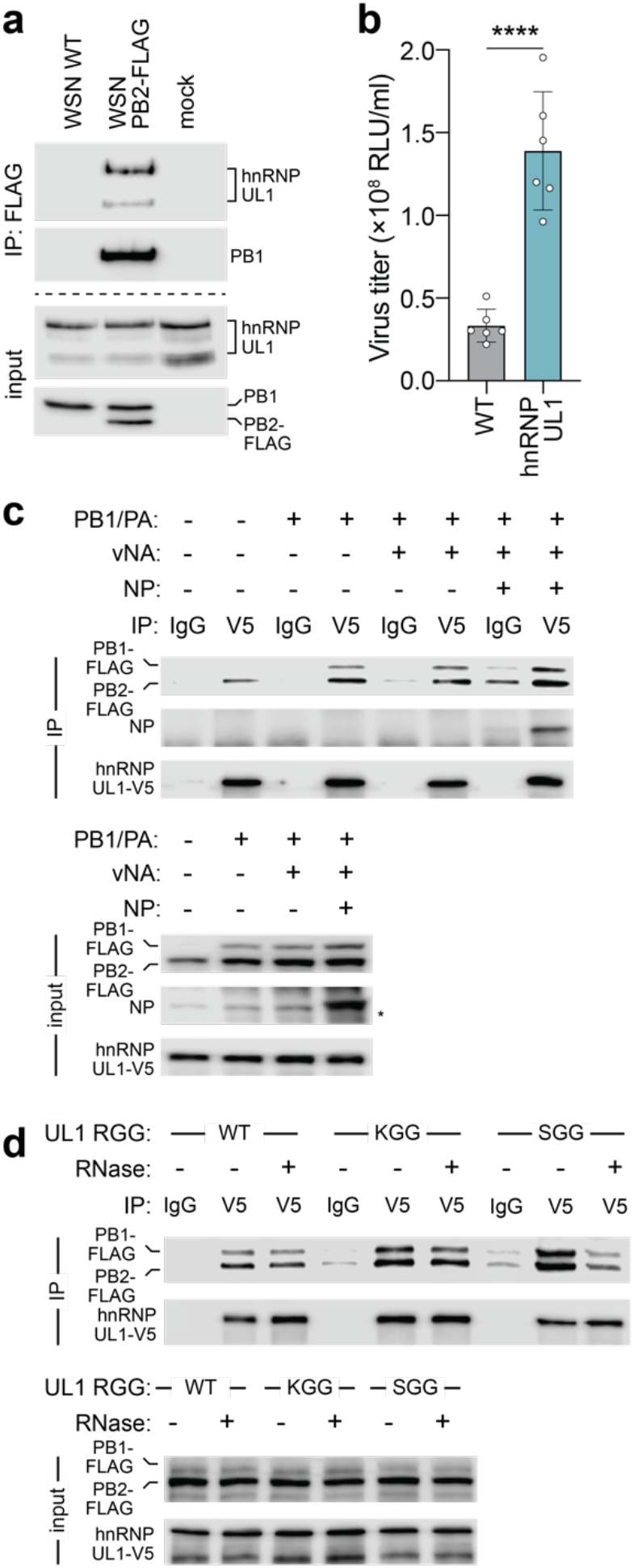
Proviral hnRNP UL1 associates with influenza polymerase. **a**, Endogenous hnRNP UL1 co-precipitates with PB2 during infection. A549 cells were infected (MOI, 1; 18 h) or mock treated, lysed and immunoprecipitated. Proteins were detected by western blot. **b**, Viral titers were measured from wildtype or clonal A549 hnRNP UL1-V5 cells infected with WSN Nluc (MOI, 0.05; 24 h). Mean ± SD of *n* = 6. Unpaired two-tailed t test; ****, *P* < 0.0001. **c-d**, Association of viral polymerase with hnRNP UL1. **c**, hnRNP UL1-V5 was immunoprecipitated with anti-V5 antibody or control IgG from lysates generated from cells expressing the indicated viral proteins or RNA (vNA).Proteins were detected by western blotting. Asterisk indicates non-specific band. **d**, Co-immunoprecipitation assays were performed using WT hnRNP UL1 or mutants disrupted in methylation (KGG) or RNA binding (SGG). RNase was included during immunoprecipitation where indicated.

Nucleocytoplasmic transport of viral mRNA is a major bottleneck for viral gene expression. There is a limiting amount of host NXF1 which chaperones mature viral mRNA to the nuclear basket for transport^33,39^. hnRNP UL1 RNP complexes with NXF1^39^. To test the importance of hnRNP UL1 and whether it is limiting, we stably over-expressed hnRNP UL1. Viral titers increased almost 3-fold in cells expressing more hnRNP UL1 (**Fig. 3b**). Similar results were demonstrated upon over-expression of NXF1 or TPR, which connects viral mRNA to the nuclear pore complex, further demonstrating that nuclear export is a limiting step during infection (**Supp. Fig. 3a**; ^40^). The viral polymerase is an RNA-binding protein that exists both in a free form or incorporated into viral RNP with genomic RNA and NP. We performed a series of interactions studies to dissect the complexes that interact with hnRNP UL1 and the role of RNA in these interactions. hnRNP UL1 interacted with PB2 when PB2 was expressed by itself (**Fig. 3c**). hnRNP UL1 showed more robust interactions when all three polymerase subunits were present, indicating that hnRNP UL1 interacts with the trimeric polymerase in the absence of other viral proteins or RNAs. Expressing genomic RNA in these cells did not increase the interaction between the polymerase and hnRNP UL1. However, including genomic RNA and NP, which permits formation of viral RNP, resulted in specific co-precipitation of NP (**Fig. 3c**). hnRNP UL1 is also an RNA-binding protein that forms multi-protein complexes. hnRNP UL1 binds RNA through its RGG box and methylation of arginine residues in the RGG box facilitates association with some of its protein partners^41,42^. To evaluate these functions, we mutated hnRNP UL1 to eliminate RNA binding (RGG to SGG) or methylation of the RGG box (RGG to KGG) (**Fig. 3d**). Interactions between the viral polymerase and hnRNP UL1 were indistinguishable between WT, an RNA-binding mutant, or a methylation mutant. These mutations did not change the normal nuclear localization of hnRNP UL1 (**Supp. Fig. 3b**). To further test for a role of bridging RNA, exogenous RNase was added during immunoprecipitation. RNase treatment resulted in minor changes for WT and The RGG, while complex formation appeared reduced for the SGG mutant, suggesting this variant may rely more on RNA bridging to engage the polymerase (**Fig. 3d**). Together, these results show that hnRNP UL1 enhances replication by binding to the viral polymerase and replication complex, potentially increasing access to rate-limiting pathways.

### *MECR* antiviral activity is independent from its role in mtFAS

MECR is an essential enzyme for mitochondrial fatty acid synthesis. The acyl carrier protein (ACP) encoded by *NDUFAB1* scaffolds each enzymatic step of mtFAS in the mitochondria with MECR performing the final step converting trans-2-enoyl-ACP to acyl-ACP (**Fig. 4a**; reviewed in ^43^). Acyl-ACP feeds into oxidative phosphorylation or is converted to octanoyl-ACP then lipoic acid for protein lipoylation or use in the TCA cycle. To test whether the mtFAS pathway itself has an antiviral role, we knocked down ACP, which potently limits lipoic acid accumulation^44^. Contrary to the antiviral activity of MECR, knockdown of ACP did not cause a significant change in viral titers (**Fig 4b**). Combined knockdown of MECR and ACP maintained the increased replication associated with MECR knockdown but did not have any additive effects. We additionally targeted mtFAS by treating infected cells with C75, a drug that targets OXSM in the mtFAS pathway as well as FASN that directs cytoplasmic fatty acid synthesis^45^. Whereas specific knockdown of MECR increased replication, complete loss of fatty acid synthesis by C75 treatment reduced viral titers, likely by disrupting cellular metabolism (**Fig. 4c**).

**Fig. 4:**
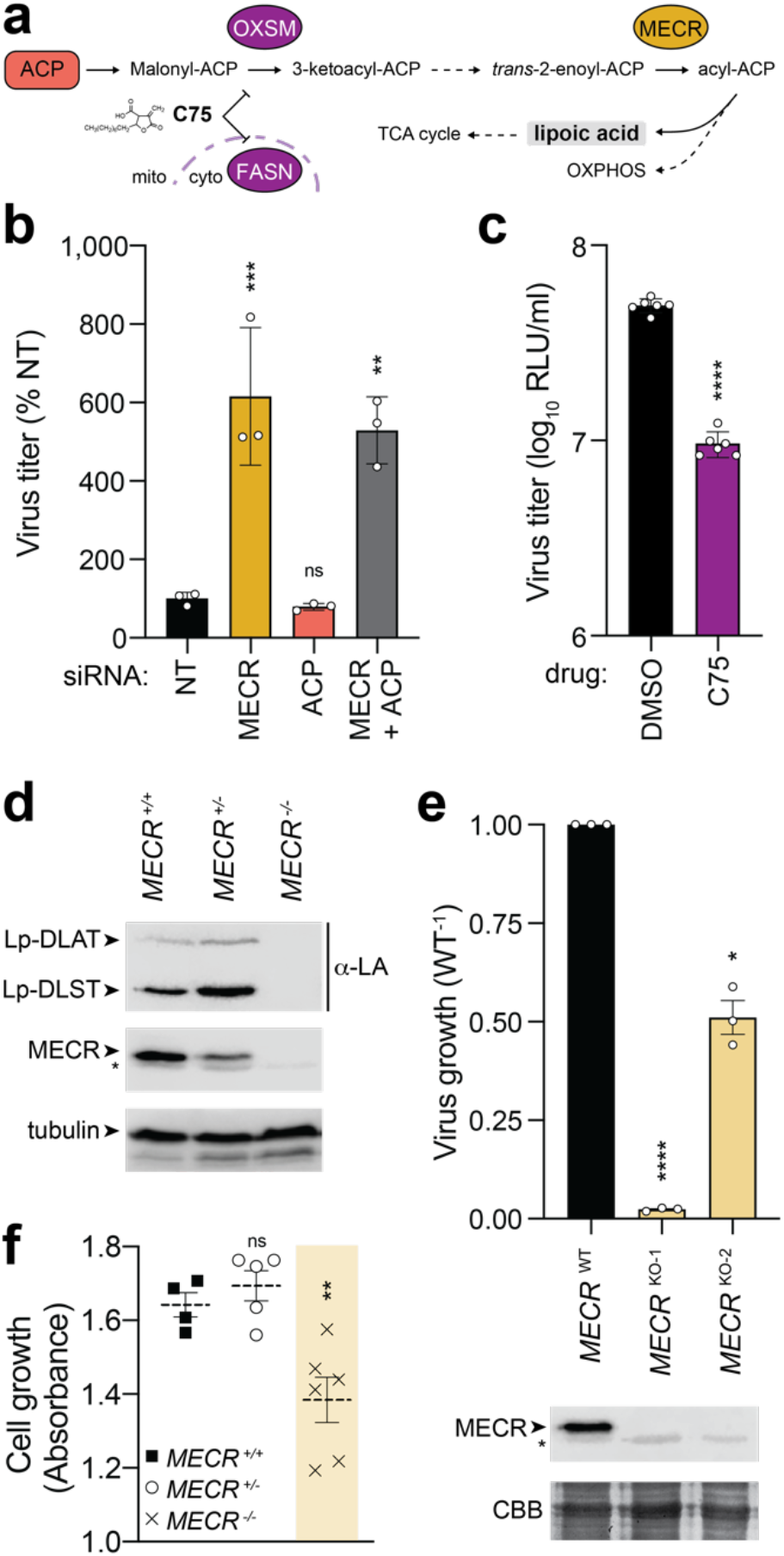
Modulating the critical mtFAS enzyme MECR alters mtFAS output and virus growth. **a**, Focused snapshot of the mtFAS pathway. Acyl-carrier protein (ACP) and MECR were experimentally probed by knockdown or knockout, whereas 3-Oxoacyl-ACP Synthase, Mitochondrial (OXSM) and fatty acid synthase (FASN) were inhibited with the drug C75. **b**, A549 cells were treated with siRNA targeting MECR, ACP, both, or a non-targeting (NT) control prior to infection with WSN NLuc virus (MOI, 0.05; 24 h). Viral titer in supernatants was determined and normalized to NT. Mean ± SD of *n* = 3. One-way ANOVA with *post hoc* Dunnett’s multiple comparisons test; **, *P* < 0.01; ***, *P* < 0.001; ns, not significant. **c**, Viral yield was measured from A549 cells treated with C75 or DMSO control prior to infection with WSN NLuc (MOI, 0.05; 24 h). Mean ± SD of *n* = 6. Unpaired two-tailed t test; ****, *P* < 0.0001. **d**, Production of lipoylated subunits of the pyruvate dehydrogenase complex (dihydrolipoamide acetyltransferase; DLAT) and the 2-oxoglutarate dehydrogenase complex (dihydrolipoamide S-succinyltransferase; DLST) was assessed in WT (^+/+^), heterozygous (^+/-^), and homozygous (^-/-^) *MECR* knockout A549 cells by western blotting with anti-lipoic acid (α-LA) antibody. **e**, Virus replication was measured in A549 cells (MOI, 0.05; 24 h). Replication in MECR knockout clones KO-1 and KO-2 was normalized to wildtype (WT) A549 cells. Mean ± SEM of biological replicates (*n* = 3) normalized to WT. MECR expression was monitored by western blotting. Coomassie brilliant blue (CBB) staining was used as a loading control. Asterisks indicate non-specific bands. **f**, Growth of clonal WT, heterozygous, or homozygous *MECR* knockout A549 cells was measured over three days. Mean ± SEM of *n* = 4-6 clones. One-way ANOVA with *post hoc* Dunnett’s multiple comparisons test; **, *P* < 0.01; ns, not significant.

To further dissect the antiviral role of MECR and any contribution from mtFAS, we generated MECR knockout A549 cells. *MECR* is not a strictly essential gene in cell culture^46^, but deletions in mice are embryonic lethal and it may be necessary for mammalian skeletal myoblasts^47,48^. We recovered A549 cells with edited heterozygotic and homozygotic knockout alleles (**Supp. Fig 4a**). Loss of MECR completely disrupted mtFAS, as indicated by loss of lipoylated proteins (**Fig. 4d**). mtFAS remained intact in a heterozygotic cell clone. Our knockdown experiments predicted that loss of MECR would enhance virus production, yet we observed that two independent MECR knockout cells uniformly produced less virus than wildtype (**Fig. 4e**). *MECR* knockout cells grew slower compared to WT or *MECR* heterozygotic cells (**Fig. 4f**). The limited virus growth in knockout cells is likely not connected to the antiviral activity of MECR, but is instead due to cellular defects in mtFAS that manifest as a loss of mitochondrial lipoic acid synthesis, slower cell growth, and defects in respiratory chain complex integrity (**Fig. 4e,f**; ^44^). These data suggest that MECR antiviral activity is independent from its normal role in supporting cellular metabolism.

### A cryptic isoform of *MECR* localizes to the cytoplasm and targets the polymerase to disrupt viral RNP assembly

Our ICC-MS results provided high confidence data demonstrating interactions between endogenous MECR and PB2 during viral infection (**Supp. Table 1**). While a minor population of PB2 localizes to the mitochondria (**Fig. 2d**; ^49^), we did not detect robust interactions between the polymerase and full-length MECR (data not shown). Analysis of RNA-seq data from influenza virus infected A549 cells revealed that 29% of MECR transcripts (± 5%, *n* = 3) have an alternative splicing pattern. The alternative transcript form utilizes an upstream splice acceptor site between exon 1 and 2 (**Fig. 5a**), which encodes multiple upstream-open reading frames and stop codons in all frames, thus creating an extensive untranslated region that may promote initiation at the downstream start site M77 in MECR. RT-PCR demonstrated the presence of long and short isoforms of MECR transcripts in mock and infected cells (**Supp. Fig. 5a-b**).

**Fig. 5:**
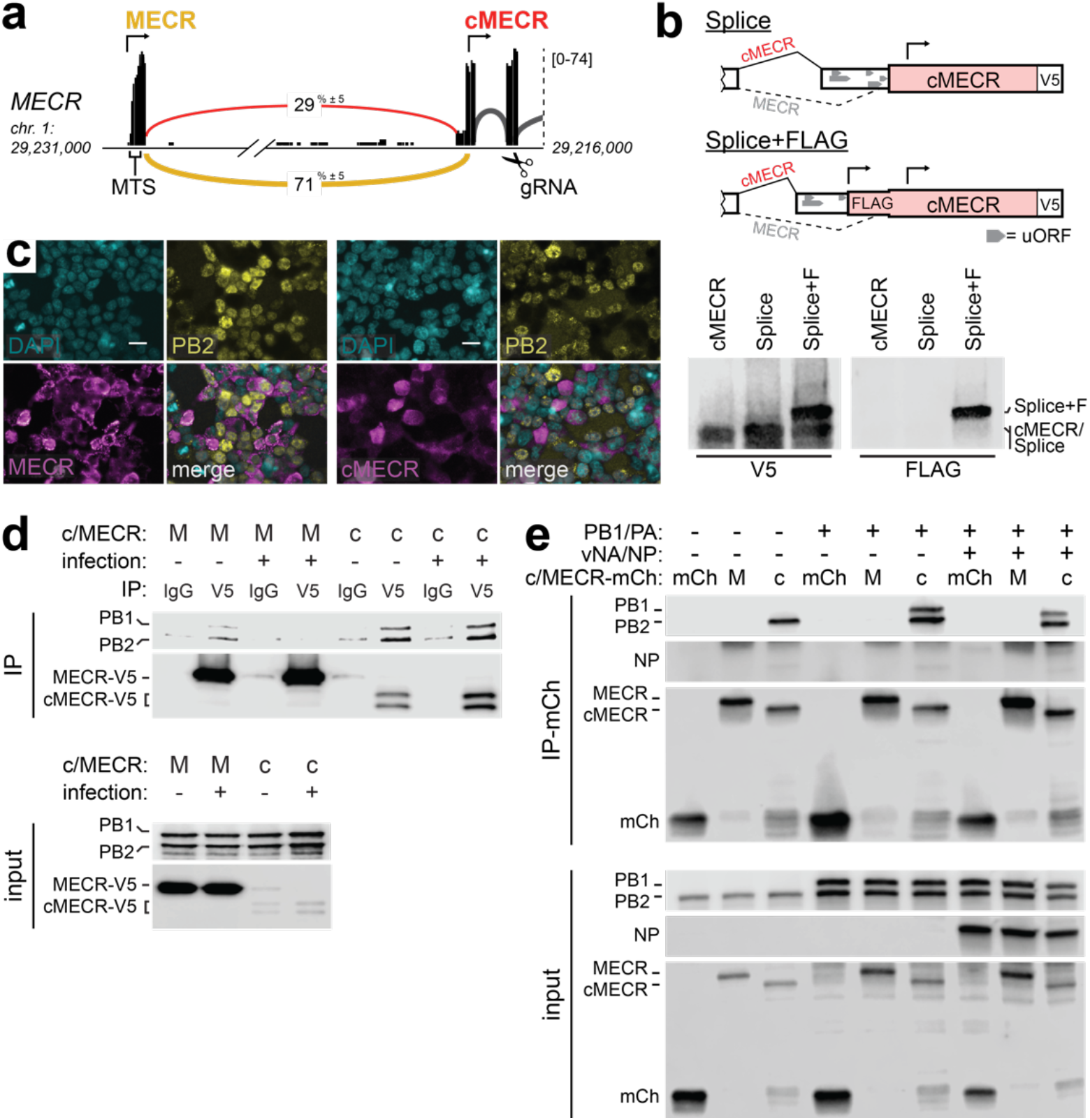
The alternative splice variant cMECR binds viral polymerase subunit PB2. **a**, Sashimi plot of RNA-seq data from A549 cells showing alternative 3’ splice site utilized by transcripts that do not translate the mitochondrial targeting signal (MTS) in exon Numbers embedded in yellow (MECR) and red (cMECR) curves indicate percentage of each splicing event as a total of all exon-joining reads. Arrows indicate translational start sites and gRNA denotes region targeted in CRISPR-Cas9 editing. **b**, Splicing reporters were constructed as diagrammed and expressed in 293T cells with cMECR cDNA as a control. Proteins were detected by western blot. **c**, 293T cells expressing MECR or cMECR were infected with influenza PB2-FLAG virus (MOI, 3; 8 h). Subcellular localization of PB2, MECR or cMECR were determined by immunofluorescence microscopy. Scale bars, 20 µm. **d-e**, The viral polymerase subunit PB2 associates with cMECR during infection. **d**, PA, PB1-FLAG, PB2-FLAG and V5-tagged MECR or cMECR were expressed in 293T cells. Where indicated, cells were also infected with WSN (MOI, 10; 6 h). Cells were lysed and immunoprecipitated with anti-V5 antibody or IgG controls. Proteins were detected by western blot. **e**, mCherry (mCh), MECR M77L-mCh, or cMECR-mCh were co-expressed with PB2-FLAG in the absence or presence of PB1-FLAG, PA, NP and viral RNA (vNA). Cells were lysed and immunoprecipitated with mCh affinity resin. Proteins were detected by western blot.

Translation from the in-frame downstream start codon would skip coding sequences for the mitochondrial targeting signal in MECR to produce cytoplasmic MECR, previously identified as cMECR^50^. Analysis of ribosome profiling data demonstrated translation initiation for cMECR at downstream sites on the alternatively spliced transcript (**Supp. Fig 5c**)^51^. Initiating ribosomes were enriched in these experiments by treating cells with lactimidomycin, a translation inhibitor that arrests ribosomes at initiation sites by blocking the first translocation cycle^52^. Under these conditions, initiating ribosomes were detected at the cMECR start site and other start sites in the UTR of the alternatively spliced transcript (**Supp. Fig. 5c**). We therefore created splicing reporters to experimentally verify translation of cMECR. The 3’ end of exon 1 and a miniaturized version of intron 1 from the *MECR* locus were cloned upstream of the minor splice acceptor site and the remainder of the MECR/cMECR cDNA (**Fig. 5b**). The reporter is designed to require splicing for cMECR production as multiple stop codons in all three frames in the intron would prevent initiation and readthrough on unspliced transcripts. The splicing reporter expressed protein similar to that from cMECR cDNA (**Fig. 5b**). To demonstrate that initiation occurs at the downstream start sites despite the presence of uORFs, we created another reporter that introduces an in-frame FLAG tag only if initiation occurs in the cMECR-specific UTR (**Fig 5b**). This reporter also produced cMECR, which is slightly larger due to the appended FLAG tag as shown by blotting. These data provide multiple lines of evidence that MECR transcripts are alternatively spliced and initiate translation at a downstream site to produce cMECR.

Full-length MECR displayed a mitochondrial-like subcellular organization, whereas expressing cMECR resulted in diffuse staining throughout the cytoplasm and nucleus, consistent with the absence of a mitochondrial targeting sequence (**Fig. 5c**). Comparing infected and uninfected cells indicated that infection did not impact MECR or cMECR distribution. Given that cMECR cannot localize to the mitochondria, it likely does not have a role in mtFAS.

As the ICC-MS results could not distinguish whether PB2 was bound by MECR or cMECR, we individually tested these interactions by co-immunoprecipitation. MECR was strongly expressed, but co-precipitated very small amounts of PB2 when PB2 was expressed alone (**Supp. Fig. 5d**). cMECR, by contrast, was poorly expressed, but robustly interacted with PB2 in the absence or presence of viral polymerase and RNP components. To evaluate if infection played a role in this interaction, parallel immunoprecipitations were performed in cells co-expressing RNP and MECR or cMECR that were additionally infected with influenza virus (**Fig. 5d**). Infection did not change the interactions; cMECR, but not MECR, interacted with the viral polymerase.

The minor co-purification of polymerase with MECR may be due to leaky scanning and initiation from M77 in MECR, recreating cMECR from the MECR transcript, which can be seen as faint lower molecular weight bands in immunoprecipitations (**Fig. 5d, Supp. Fig 5d)**. We introduced an M77L mutation in *MECR* to overcome leaky initiation. We also codon-optimized constructs to improve cMECR expression and fused them to mCherry for visualization. These changes did not alter the subcellular localization of the expressed proteins (**Supp. Fig. 5e**). Immunoprecipitations were repeated with these new constructs that strictly express MECR or cMECR. cMECR, but not MECR, again co-precipitated PB2 when it was expressed alone and within the context of the trimeric viral polymerase (**Fig. 5e**). cMECR also co-precipitated PB1 when it was present, although NP was notably absent from these interactions. We utilized proximity ligation as an orthogonal approach to query interactions with cMECR. Polymerase was co-expressed with cMECR fused to the biotin ligase AirID^53^. In the presence of cMECR-AirID, purified polymerase was heavily biotinylated on PB2 and PA, and to a lesser extent on PB1 (**Supp. Fig. 5f**). These targeted interaction studies, combined with endogenous interactions detected by ICC-MS, provide multiple lines of evidence that PB2 interacts exclusively with cMECR.

Given that cMECR interacted with the viral polymerase, we asked whether it conferred the antiviral phenotype revealed by our knockdown experiments (**Fig. 2a, c**). Viral titers were measured following infection of clonal A549 cells expressing MECR or cMECR. Overexpression of MECR did not alter virus replication, whereas exogenous cMECR resulted in a nearly 10-fold reduction in viral titers (**Fig. 6a**). Successful infection requires *de novo* formation of viral RNPs to amplify transcription and replicate the viral genome. cMECR co-precipitated polymerase in cells where RNPs are formed, yet NP was not co-precipitated, suggesting cMECR interacted with free polymerase but not polymerase assembled into RNPs (**Fig. 5e**). We thus tested whether cMECR interferes with RNP assembly (**Fig. 6b**). Lysates were prepared from cells where RNPs were assembled in the presence of MECR M77L, cMECR or the control mCherry. The lysates were split and fractions were used to assess interactions with cMECR or RNPs. In one fraction, cMECR again co-precipitated the viral polymerase, but not NP. MECR and the mCherry control did not interact with viral proteins, confirming specificity. In the other fraction, NP co-precipitated the polymerase, confirming RNP formation. However, expressing cMECR reduced RNP formation compared to MECR or the mCherry control. Moreover, the NP-captured RNP complexes were devoid of cMECR. To further test the ability of cMECR to disrupt RNP formation, reciprocal immuno-precipitations were performed where the viral polymerase was captured and probed for co-precipitating NP (**Fig 6c**). cMECR again reduced RNP formation as seen by lower amounts of NP interacting with the polymerase compared to MECR. These complementary analyses suggest mutually exclusive protein complexes where cMECR interacts with free polymerase and prevents its incorporation into RNPs.

**Fig. 6:**
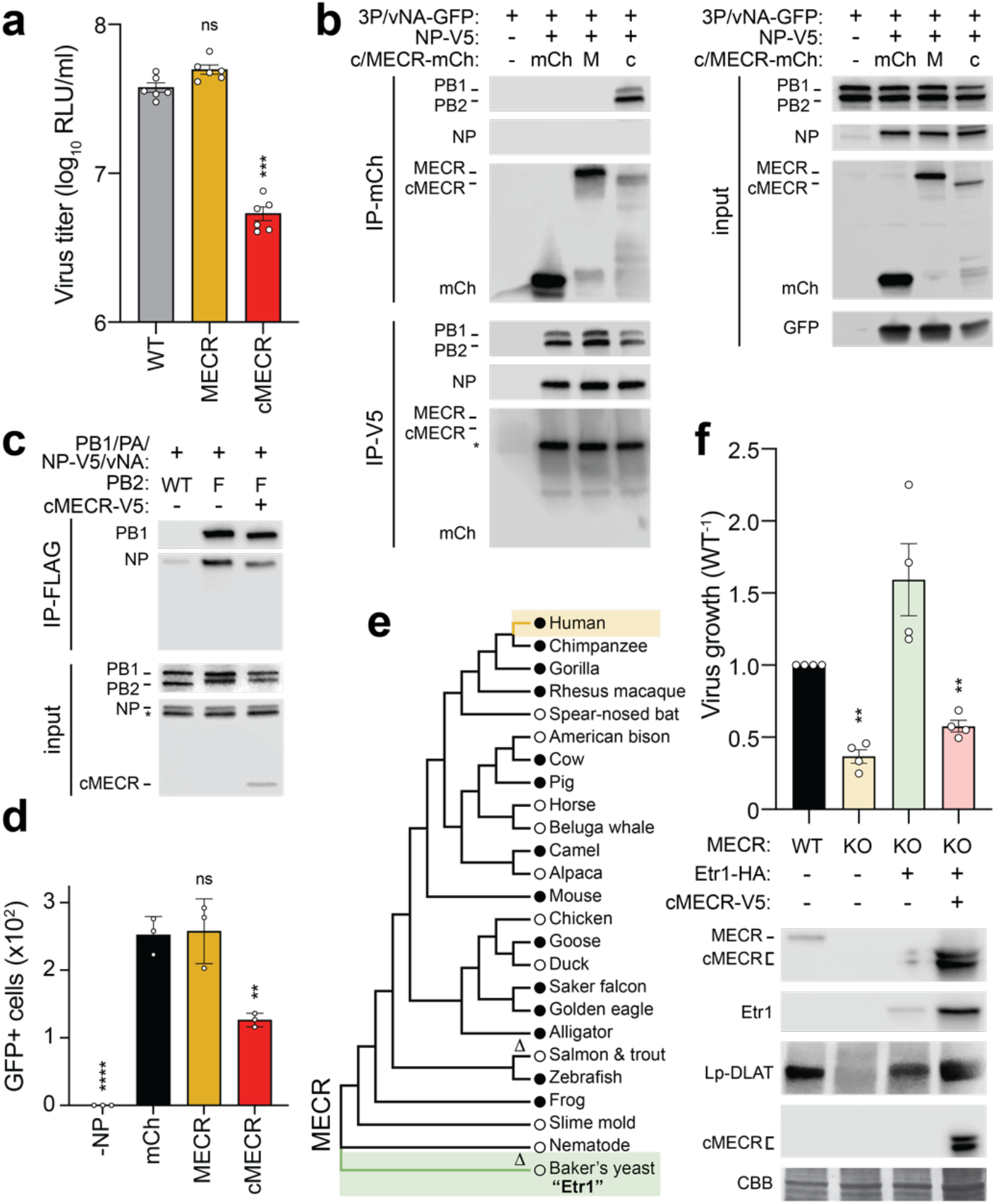
The antiviral activity of cMECR inhibits RNP assembly independent of MECR and mtFAS. **a**, Replication of WSN NLuc was measured in WT or clonal A549 cells expressing MECR or cMECR. Protein expression from infected cells was analyzed by western blotting. Asterisk indicates a non-specific band used as a loading control. Mean ± SD of *n* = 6. One-way ANOVA with *post hoc* Dunnett’s multiple comparisons test; ***, *P* < 0.001; ns, not significant. **b-c**, cMECR disrupts RNP assembly. **b**, RNPs were assembled in 293T cells co-expressing mCh, MECR M77L-mCh, or cMECR-mCh. NP was omitted in the negative control (-NP). Cells were lysed, divided in half, and immunoprecipitated for MECR (mCh) or NP (V5). Input or co-precipitated proteins were detected by western blot. * = NP detected from prior probing of the membrane before blotting for mCh. **c**, RNP assembly monitored as in b, except PB2 (FLAG) was targeted for immunoprecipitation with untagged PB2 as the negative control. **d**, Polymerase activity assays with vNA-GFP were performed in the presence of mCh, MECR M77L-mCh, or cMECR-mCh. Fluorescence microscopy images taken at 24 h post-transfection and GFP-positive cells were enumerated by Image J. Mean ± SD of *n* = 3. Two-way ANOVA with *post hoc* Dunnett’s multiple comparisons test; ***, *P* < 0.001; **, *P* < 0.01; ns, not significant. **e**, Phylogenetic maximum-likelihood analysis of MECR amino acid sequences. Δ, sequences lacking conserved cMECR start codon (Met77, human numbering). Circles indicate presence (filled circles) or absence (empty circles) of annotated cMECR transcripts. NCBI RefSeq IDs listed in Supp. Table 4. **f**, MECR KO cells were complemented with the MECR homolog from *S. cerevisiae* Etr1 then transduced with cMECR. Cells were subsequently infected with WSN NLuc (MOI, 0.05; 24 h) and viral titers were measured in supernatants and compared to WT. Mean ± SD of *n* = 4. Unpaired two-tailed t test; *, *P* < 0.05; **, *P* < 0.01. *below*, mtFAS rescue was confirmed by detecting Lp-DLAT via western blotting whole cell lysates.

If cMECR interferes with RNP assembly, it should also decrease polymerase activity. RNPs assembled in the above experiment contained a model vRNA encoding GFP. Polymerase activity was decreased in cells expressing cMECR, as seen by a reduction in GFP protein levels compared to cells expressing MECR M77L or the control mCherry (**Fig. 6b, input)**. Co-expressing cMECR also caused a ∼50% decrease in GFP-positive cells compared to the mCherry control, while full length MECR M77L had no appreciable effect (**Fig. 6d**; **Supp. Fig. 6a**). Collectively, these data suggest that cMECR exerts its antiviral activity by suppressing RNP assembly and downstream polymerase activity.

### Repairing mtFAS with the yeast homolog assigns antiviral activity to cMECR

MECR plays an essential role in mtFAS, making it challenging to cleanly delineate MECR function during mtFAS from the antiviral role of cMECR. Indeed, mtFAS deficiency in *MECR*^*-/-*^ cells resulted in a generic defect in cellular metabolism that reduced cell growth, and consequently viral replication (**Fig. 4**). We therefore sought to repair the mtFAS pathway in *MECR*^*-/-*^ cells by expressing a homolog of MECR that lacks cMECR. cMECR translation relies on the start codon at M77, which is surprisingly well-conserved; phylogenetic analysis suggests that the M77 start codon driving cMECR expression was acquired after the divergence of animalia from fungi (**Fig. 6e**). Only the ancestral homolog of MECR from baker’s yeast (*Saccharomyces cerevisiae*) and salmon (*Salmo salar*) and trout (*Salmo trutta*) do not encode M77 and presumably do not produce cMECR. cMECR expression also requires an alternatively spliced transcript. Annotated transcripts that code for cMECR were identified across many influenza virus hosts including pigs, mice, and geese, providing further evidence for cMECR expression (**Fig. 6e, Supp. Table 4**).

To assess whether the antiviral activity of cMECR is independent from mtFAS and MECR in general, we complemented *MECR*^*-/-*^ cells by stably expressing the yeast homolog Etr1. As before, knockout of MECR disrupted mtFAS as shown by the loss of lipoylated DLAT and reduced viral replication compared to WT cells (**Fig. 6f**). Etr1 functioned in human cells to restore lipoic acid synthesis indicated by the return of lipoylated DLAT in the complemented cells. Furthermore, Etr1 expression also increased virus growth to greater than WT levels, likely due to the absence of cMECR. We then used the Etr1-complemented genetic background to demonstrate that cMECR alone impairs virus growth (**Fig. 6f**). These results were confirmed in two independent knockout cell lines (**Supp. Fig. 6b**). The genetic complementation approach also demonstrated that the anti-influenza activity of cMECR does not require the presence of full-length MECR, which is important considering MECR is known to homodimerize^54^. Collectively, our results decouple the newly described antiviral activity of cMECR from the canonical role of MECR.

## Discussion

Zoonotic influenza viruses must gain a foothold to replicate and then adapt to their new human hosts. Here we used ICC-MS to identify human proteins that interact with influenza polymerases containing avian or human-signature PB2 during infection. Network simulations and siRNA screening highlighted two key interactors, hnRNP UL1 and cMECR. We showed that hnRNP UL1 supports viral replication and likely acts as part of a pathway commonly exploited by viruses to aid mRNA splicing and transport. cMECR, by contrast, exerted antiviral activity by inhibiting RNP assembly and thus suppressing viral gene expression and replication. MECR has a well-defined role in mitochondrial fatty acid synthesis, yet this pathway was not identified in our network analyses and mechanistic studies showed that the antiviral activity was independent of mtFAS. MECR and mtFAS were generically important for viral replication, in as much as they were necessary for proper mitochondrial metabolism and cell health. Instead, we demonstrate the antiviral activity derives from cMECR, an alternative splice variant that lacks the mitochondrial targeting sequence and is cytoplasmic. Using the yeast homolog Etr1 to supply the metabolic functions of MECR, we confirmed that antiviral activity is independent of mtFAS and lies solely within cMECR. Thus, a cryptic antiviral activity has been embedded within a key metabolic enzyme, possibly protecting it from viral antagonism.

hnRNP family members, of which there are 37 in humans, broadly regulate host nucleic acid processes including chromatin organization, DNA damage repair, pre-mRNA processing and splicing, and mRNA nuclear export and subcellular transport^55,56^. As a consequence, hnRNPs are frequent co-factors for viral replication, including hnRNP UL1 we characterized here^57^. hnRNP UL1 contains RGG domains that can be methylated and are involved in RNA binding. However, neither of these activities were required for interaction with the viral polymerase (**Fig. 3e**). hnRNP UL1 also binds to NXF1, the major cellular factor involved in nuclear export of viral mRNAs^33,39,58^. We speculate that hnRNP UL1 may enhance viral replication by bridging interactions between transcribing viral polymerases and NXF1 to facilitate mRNA export.

Mitochondria are a major subcellular location where host-pathogen conflicts play out: cellular IFN pathways use the mitochondrial membrane to assemble signaling hubs containing MAVS; mitochondrial nucleic acids can be unshielded to trigger the type-I IFN response; and, miRNAs that regulate IFN stimulated genes are embedded in mitochondrial genes^59–62^. However, antiviral activity of *MECR* does not require innate sensing pathways that converge at the mitochondria as knockdown of *MECR* in A549 MAVS, RIG-I, or PKR knockout cells still alleviates its antiviral activity (**Supp. Fig. 6c**). Instead, our data uncover another conserved mechanism where the metabolic enzyme MECR uses mRNA alternative splicing, and possibly leaky scanning, to moonlight as the antiviral protein cMECR. cMECR limits influenza A and B viruses, but not pandemic 2009 H1N1 (CA04) (**Fig. 2e**). WSN and B/Bris are human origin strains, whereas the PB2 gene from CA04 is avian-like^63^. Human-derived PB2 genes localize to the mitochondrial matrix due to a mitochondrial targeting sequence in the N-terminus; the targeting sequences are disrupted in avian PB2 proteins by an N9D polymorphism^49,64,65^. Whether this contributes to differences in sensitivity to cMECR remains to be explored.

Phylogenetic analysis revealed that both MECR and the M77 that initiates cMECR are highly conserved in typical influenza hosts, including humans, birds, pigs, and horses. It is not yet known whether the absence of annotated cMECR transcripts in some species is due to *bona fide* differences in splicing or failed prediction algorithms. Amongst this conservation, the loss of M77 and potentially cMECR in *Salmo* species is especially interesting (**Fig. 6e**; human amino acid numbering). *Salmo* species are hosts for infectious salmon anemia virus, an *orthomyxovirus* closely related to influenza virus. Analysis of splicing events in RNA-Seq data shows that *Salmo* do not appear to utilize the alternative splice acceptor site that could create cMECR. Nonetheless, they may still express a cMECR-like protein by initiating translation at a different start codon, possibly the conserved M89 of MECR. This would require a distinct mechanism, such as leaky scanning, rather than the alternative splicing used for cMECR production in other vertebrates. *Salmo* do vary splicing between exon 1 and 2 by utilizing an alternative splice donor to create a splice isoform that encodes 26 amino acids not found in humans. It will be important to determine if *Salmo MECR* encodes a protein with antiviral activity, providing information on the antiviral function of MECR in different species and illuminating how conditions that alter splicing patterns or translation initiation sites stimulate or repress cMECR production.

Viruses often exploit conserved essential host genes whose critical role for the cell limits mutational escape. Conversely, host antiviral genes are often nonessential and can undergo mutation or regulated expression to thwart infection and limit self damage. Our study provides an example of a gene that is both essential and antiviral. Embedding the antiviral cMECR within the essential metabolic enzyme MECR establishes a Corneillian dilemma for influenza virus. Viral antagonism of cMECR could have the off-target effect of disabling MECR and mtFAS, a metabolic process important for cell viability and influenza virus output. Conversely, leaving MECR and mtFAS intact would ensure cell health, while the virus would remain vulnerable to cMECR. The host strategy of encoding an alternatively spliced antiviral protein that shares structure with an energetically important protein may represent a failsafe for the host, forcing the virus to make the difficult “decision” of antiviral antagonism or maximum biosynthetic power.

## Acknowledgements

We thank members of the Mehle lab for critical reading of the manuscript. We thank Craig McCormick (Dalhousie University) and Nathan Sherer (University of Wisconsin-Madison) for sharing reagents. This work was supported by NIH/NIAID R01AI164690 and the Greater Milwaukee Foundation Shaw Scientist Award to AM, R21AI125897 to AM and AG, a Roche Postdoctoral Fellowship RPF-353 and the NIAID T32AI55397 to SFB, T32LM012413 to AB, and an NSF GRFP DGE-1747503 to MPL. AM is a Burroughs Wellcome Fund Investigator in the Pathogenesis of Infectious Disease and an H. I. Romnes Faculty Fellow funded by the Wisconsin Alumni Research Foundation.

## Author contributions

Conceptualization: SFB, HM, MT, AB, AG, AA, HJ, AM

Methodology: SFB, HM, MT, AB, SG, AA, AM

Formal Analysis: SFB, HM, MT, AB, JSP, MPL, AG, AA, AM

Investigation: SFB, HM, MT, AB, MPL, AM

Writing – Original Draft: SFB, AB, AG, AM

Writing – Review & Editing: all

Visualization: SFB, AB, AM

Funding Acquisition: SFB, MPL, AB, AG, AM, HJ

Supervision: HM, AG, AA, HJ, AM

## Methods

### Viruses, cells, plasmids, antibodies

Influenza viruses and plasmids were derived from A/WSN/33 (H1N1; WSN), A/green-winged teal/Ohio/175/1986 (H2N1; S009), A/California/04/2009 (H1N1; CA04), and B/Brisbane/60/2008 (B-Victoria lineage; B/Bris)^18,28,66,67^. Recombinant virus was rescued by transfecting co-cultures of 293T and MDCK cells with pTM RNP encoding WSN vRNA segments HA, NA, M, and NS and the bi-directional pBD plasmids encoding vRNA and mRNA for PB1, PA, NP, and the indicated PB2 mutants^18,68^. WSN PB2-FLAG and WSN-PB2-627E-FLAG virus were previously described^35,69^. S009 PB2-FLAG and S009 PB2-627K-FLAG viruses contain NP and polymerase genes from S009 and the remaining segments from WSN and were constructed as previously described^18,70^. Nanoluciferase-expressing (NLuc) viruses were engineered to co-linearly express PA, a 2A cleavage site, and NLuc (PASTN, referred to as WSN NLuc) on the third viral segment^71^. CA04 NLuc and B/Bris NLuc were generated similarly^28,67^. Viral stocks were amplified on MDBK cells and titered by plaque assay on MDCK cells. Influenza virus infections were performed by inoculating cells with stocks diluted in virus growth media (VGM; DMEM supplemented with penicillin/streptomycin, 25 mM HEPES, 0.3% BSA, and 0.25 – 0.5 µg/ml TPCK-trypsin).

VSV-G pseudotyped lentivirus was prepared by transfecting 293T cells with pMD2.G (Addgene 12259), pLX304 (Addgene 25890) encoding HNRNPUL-1-V5 or MECR-V5; HNRNPUL1-V5 and MECR-V5) and psPAX2 (Addgene 12260) or pMD2.G, pQCXIP encoding Etr1-HA (Clontech) and pCIG-B^72^. Resultant viruses were used to transduce A549 cells. Cells were selected with blasticidin or puromycin to obtain stable expressing lines.

Mammalian cells were grown in DMEM supplemented with 10% FBS: 293T, ♀; A549, ♂; MDBK, ♂; MDCK, ♀. Innate sensing A549 knockout cells were a kind gift from C. McCormick and described previously^73^. All cells were maintained at 37 °C, 5% CO_2_. Cell stocks were routinely tested for mycoplasma (MycoAlert, Lonza). Human cell lines were authenticated by STR analysis (University of Arizona Genetics Core).

A549 MECR knockout cells were generated using CRISPR-Cas9 with a single guide RNA targeting exon 3 of *MECR* (GTTGCACAGGTGGTAGCGGTGGG designed by crispr.mit.edu). Annealed tracrRNA/gRNA was complexed with Cas9 (Alt-R; IDT) and RNPs were electroporated (Lonza) into cells following manufacturer’s instructions. Two days post nucleofection, bulk population (42% editing efficiency by sanger sequencing and ICE analysis^74^) was single cell sorted into 96 well plates by FACS. Clones were screened by targeted next generation sequencing of genomic DNA (Genome Engineering & iPSC Center, Washington University in St. Louis). Biallelic knockout cells were verified by western blot.

Cell proliferation assays were conducted over three days using CellTiter 96 AQ reagent (Promega) at 24 or 72 h after seeding, incubating 2 h, and measuring absorbance. Background absorbance at 630 nm was subtracted from 490 nm. Measurements were performed in technical triplicate and activity from 72 h subtracted from 24 h to determine growth.

pENTR HNRNPUL1 was generated using Gibson assembly from pDONR HNRNPUL1 obtained from DNASU.org (HsCD00719283). RNA binding mutants of HNRNPUL1 were generated by synthesizing mutant RGG boxes (nucleotides encoding amino acids 612-658; IDT) where all arginines were replaced with serine (SGG) or lysine (KGG) and introduced using Gibson assembly^41,42^. pDONR MECR was acquired from DNASU (HsCD00399762). pDONR cMECR was generated using inverse PCR to delete nucleotides encoding the first 76 amino acids of MECR. HNRNPUL1 and MECR were recombined into V5-tagged mammalian expression constructs by Gateway cloning into pcDNA6.2 (Invitrogen) and pLX304 (Addgene 25890). Codon optimized MECR was synthesized (IDT) containing an M77L mutation and cloned into pcDNA3 fused to a GGSGG (5GS) linker and mCherry (mCh) coding sequence on the 3’ end. Codon optimized cMECR was generated by inverse PCR using codon optimized MECR as a template. pQCXIP (Qiagen) Etr1-HA was generated by Gibson assembly using pDONR Etr1 (DNASU; ScCD00009122). pcDNA3 FLAG-TPR was acquired from Addgene (60882). pcDNA6.2 NXF1 was generated by Gateway cloning pENTR NXF1 (DNASU; HsCD00514182).

Antibodies used for blotting include monoclonal anti-FLAG clones 1/27 and 1/54 made in-house at Roche, anti-FLAG M2-HRP (Sigma A8592), polyclonal anti-MECR (Proteintech 14932-1-AP or Atlas Antibodies HPA028740), polyclonal anti-hnRNP UL1 (Proteintech 10578-1-AP), polyclonal anti-V5 (Bethyl Labs A190-120A), monoclonal anti-V5-HRP (clone V5-10, Sigma V2260), polyclonal anti-PB1^18^; monoclonal anti-tubulin (clone DM1A, Sigma T6199); polyclonal anti-lipoic acid (Calbiochem 437695); monoclonal anti-HA-HRP (clone 3F10, Sigma 12013819001), polyclonal anti-rabbit-HRP (Sigma A0545); and rabbit IgG (2729, Cell Signaling Technology). Antibodies used for immunofluorescence include mouse monoclonal anti-FLAG (Sigma F1804), rabbit polyclonal anti-V5 (Bethyl Labs A190-120A), goat anti-mouse Alexa Fluor 594 (Invitrogen A-11032), goat anti-rabbit Alexa Fluor 488 (Invitrogen A-11008).

### Immuno-competitive capture (ICC)

ICC was performed as previously published^17^ with the following modifications. Anti-FLAG clone 1/27 was coupled to Affi-Gel 10 resin following the manufacturer’s instructions (Bio-Rad). A549 cells were inoculated at an MOI of 0.2 with S009 PB2-FLAG or S009 PB2-627K-FLAG in a 10 cm dish, 5 dishes per replicate, 3 biological replicates. Infections were allowed to proceed for 24 h. Cells were combined and lysed in 1 ml co-IP buffer (50 mM Tris, pH 7.4, 150 mM NaCl, 0.5% NP-40, 1X cOmplete protease inhibitor (Roche)) and divided into equivalent fractions for ICC. Lysates were incubated by rocking for 3 h at 4 °C with free competing antibody (anti-FLAG 1/27) where applicable, then immunoprecipitated for 16 h with 1/27 antibody coupled to Affi-Gel 10 resin. After immunoprecipitation, samples were washed four times with co-IP buffer and eluted in Laemmli buffer at 70 °C for 10 min. Samples were then transferred to new tubes and boiled for 10 min with 0.1 M DTT. 10% of the immunoprecipitates were separated by SDS-PAGE and analyzed by western blotting using 1/54 anti-FLAG antibody or silver stained.

### Mass spectrometry (MS)

The remaining 90% of the ICC sample was separated on a 4-20% Tris-Glycine SDS-PAGE gel and stained with Coomassie blue. Lanes were cut from the gel and processed for in-gel digestion. Samples were analyzed with a nanoflow Easy-nLC 1000 system (Proxeon) connected to an Orbitrap Fusion Tribrid mass spectrometer and equipped with an Easy-spray source (Thermo Fisher Scientific). Samples were re-suspended in LC-MS buffer (5% formic acid/2% acetonitrile), concentrated on an Acclaim PepMap C18 trapping column (75 *µ*m × 20 mm, 5 *µ*m particle size), and peptides separated on an Acclaim PepMap C18 EASY-spray column (75 *µ*m × 500 mm, 2 *µ*m particle size) heated at 45 °C using the following gradient at 300 nL/min: 7-50% B in 45 min, 50-80% B in 2 min, 80% B for 13 min (buffer A: 0.1% formic acid; buffer B: 0.1% formic acid/acetonitrile). The instrument was set to collect Orbitrap MS1 scans over a mass range from *m/z* 300 to 1500 using quadrupole isolation, a resolution of 120 000 (at *m/z* 200), an automatic gain control (AGC) target value of 2 × 10^5^, and a maximum injection time (IT) of 100 ms. Using data-dependent acquisition (DDA) with a cycle time of 3 s between two MS1 scans, the most intense precursor ions with a minimum intensity of 5 × 10^3^, were mono-isotopically selected for high-energy collision dissociation (HCD) using a quadrupole isolation window of 0.7 Th, AGC target of 1 × 10^4^, maximum IT of 35 ms, collision energy of 30%, and ion trap readout with rapid scan rate. Charge states between 2 and 6 and only one per precursor was selected for MS2. Already interrogated precursor ions were dynamically excluded for 20 s using a ±10 ppm mass tolerance.

MS raw files were processed for label free quantification using Progenesis QI 2.1 (Nonlinear Dynamics) and ions m/z values were aligned to compensate for drifts in retention time between runs (maximum charge state set at +5). Peptides and proteins were identified by searching data with Mascot Server 2.5.1 (Matrix Science) together with the UniProt human (May 2016 release, 20,201 sequences) and the influenza S009 viral (12 sequences) protein databases. Searches used trypsin/P as an enzyme, a maximum of two missed cleavage sites, and 10 ppm and 0.5 Da as the precursor and fragment ion tolerances, respectively. Carbamidomethylated cysteines (+57.02146 Da) were set as static while oxidized methionines (+15.99492 Da) were set as dynamic modifications. The specFDR was restricted to 1% by performing a target-decoy search using a concatenated decoy database. Peptide extracted ion chromatograms (EICs) were used to determine peptide amounts. Data normalization was performed in Progenesis by applying a scalar multiple to each feature abundance measurement with the assumption that most peptide ions do not change in abundance (similar abundance distributions globally.

Data were quality controlled by assessing sample distribution and performing a principal component analysis. One outlier sample (Dose 0) was removed from the PB2-627K experiment, based on Mahalanobis distance of the first 3 principal components^75^. Specific interacting proteins should decrease in relative abundance with increased concentration of the free competitor antibody; these displaced proteins are determined as previously described^76^. Briefly, a linear model is fit on the log_2_-transformed relative abundance values for each protein with the free competitor compound concentration. Then, monotonic contrasts are used to compare the protein abundance values above and below each concentration point^77,78^. The maximum t-statistic from each series is determined with a moderated t-test^79^. Model fitting and post-hoc contrast tests are performed with limma^80^. Significance of the displacement is assessed using permutation tests, where concentration labels were permuted 1000 times based on the step-down minP algorithm^81^ modified for one-sided tests, and adjusted for multiple testing^82^. Proteins with adjusted p-values below 5% are considered specific binders. Computations were performed in R^83^.

### Network Analysis

Network analysis was formulated as a minimum-cost flow optimization problem inspired by ResponseNet but customized for our application^84^. Human protein-protein interactions were obtained from the STRING database (version 10.5), using only interactions supported by experimental evidence^22^. PB2 interactors identified by ICC-MS were designated as source nodes and the influenza host factors as target nodes in the STRING network. The host factors came from our prior genetic screens^28,29^ and published RNA interference screens^23–27^. All human gene symbols were mapped to Ensembl protein identifiers. Proteins not present in this version of the STRING interaction network (i.e. PB2 interactors ABLIM1, EPDR1, and some influenza host factors) and TMEM106B were not included in the network analysis, leaving 23 source nodes and 2,179 target nodes.

The goal in the minimum-cost flow problem is to transport units of flow from a source node *S* in a network to a target node *T*^84^. *S* and *T* are special nodes that are added to the network and do not represent proteins. *S* has outgoing edges to all of the protein source nodes, the PB2 interactors. *T* has incoming edges from all the protein target nodes, the host factors. Flow can move from node to node through edges in the network. The total amount of flow to transport from *S* to *T* is fixed. However, each edge has its own cost associated with transporting a unit of flow over that edge and a capacity that limits how much flow it can transport. Flow was assigned to edges such that the total cost of transporting the fixed amount of flow is minimized. Additional constraints require that the total flow into a protein node in the network (that is, all nodes except *S* and *T*) equals the total flow out of that node, the total flow from *S* equals the total flow into *T*, and the flow assigned to each edge is non-negative and less than or equal to the edge’s capacity. Solving the minimum-cost flow problem assigns how much flow each edge transports. The edges with positive flow compose a subnetwork, and in our application that subnetwork may comprise a predicted host influenza response pathway.

The standard minimum-cost flow problem was adjusted here to ensure that not all flow can be transported through one source or one target and a minimum number of sources and minimum number of targets that must transport positive flow was specified. The total flow to transport is then the product of these minimums. The network edges from *S* to the sources had capacity equal to the minimum number of targets. The network edges from the targets to *T* had capacity equal to the minimum number of sources. The protein-protein interaction edges had capacity equal to the total amount of flow, which did not constrain how much flow could be transported. The protein-protein interaction edges also had costs derived from STRING. The cost was one minus the STRING weight for the interaction, such that low confidence edges have higher costs. This construction guarantees that if a feasible solution exists, it will include at least as many sources and targets as requested. Furthermore, the solution will use the most confident protein interaction edges to transport that flow. If multiple equally-good solutions exist, the solver selected one arbitrarily. If a solution could not be found, we reduced the minimum number of sources and targets. For the influenza network analysis, the minimum number of sources was set to 23 and the minimum number of targets to 200. This number of targets produced subnetworks that had multiple PB2 interactors in each connected component, which we also refer to as modules, as opposed to subnetworks that placed each source in its own connected component. The SimpleMinCostFlow solver from the ortools Python package (version 6.10.6025, https://developers.google.com/optimization) was used to solve the minimum-cost flow instances. Our Python code is available from https://github.com/gitter-lab/influenza-pb2.

Flow simulations can be biased due to nodes forming spurious links in order to reach large multi-partner targets. Two control analyses were conducted to control for this possibility and assess the significance of the protein interaction subnetwork from the minimum-cost flow solution. Both controls solve the minimum-cost flow problem many times using randomized input data that is not relevant to influenza A virus. If a protein belongs to both the influenza virus subnetwork and these control subnetworks, it may have been selected due to properties of the STRING network rather than influenza relevance. We defined an empirical *P* value for each node in the influenza subnetwork as the number of times that node appears in a control subnetwork divided by the number of control runs. We executed 1,000 control runs. The simulated control sampled 23 source nodes, the number of real PB2 interactors, uniformly at random from all nodes in the STRING network. It used the real influenza host factors as target nodes. Results demonstrated that most of the flow subnetworks containing ICC-MS hits were specific to those generated by PB2 interactors and not randomly sampled proteins (**Supp. Table 2a**). The second type of control uses an alternative virus, hepatitis C virus, as the input. The alternative control sampled 23 source nodes from the 1,864 human proteins that interact with the hepatitis C virus nonstructural protein 5A as sources^17^. It used hepatitis C host factors from a CRISPR screen as targets^85^. This simulation demonstrated that the connection of most PB2 interactors to important viral cofactor nodes was specific to the influenza virus-defined network, and not the HCV-defined network (**Supp. Table 2b**). Both controls confirm that our PB2 interactors connect to subnetworks relevant for influenza virus, and not general virus replication or generic hubs.

The node sizes in the influenza subnetwork visualizations were scaled to be proportional to the node’s negative log_10_ *P* value from the simulated and alternative controls. If a node’s empirical *P* value was 0, we set it to 0.001 for this visualization. In addition, subnetwork regions were annotated with enriched GO terms using the gprofiler-official Python package (version 0.3.5) for gene set enrichment analysis on each connected component of the subnetwork^86^. Each connected component-GO term enrichment result was ranked by a score that incorporated the negative log_10_ *P* value of the enrichment, the depth of the GO term in the biological process ontology, and the fraction of nodes in the connected component annotated with the GO term. The combined score emphasizes more specific GO terms that have statistically significant enrichment and cover a large fraction of nodes. For each connected component, we assigned the GO term with the largest combined score that had not already been assigned to a different connected component. The influenza subnetworks were visualized with Graphviz^87^.

### siRNA screening and infection experiments

A549 or 293T cells were reverse transfected with 25 nM siRNA (SMARTpool, Horizon) using siQuest or X2 (Mirus) in 96-well plates for 48 h. Cells were inoculated with PASTN virus at an MOI of 0.1 (single cycle infection, 8 h) or 0.05 (multicycle infection, 24 h). Single cycle monolayers were seeded in white bottom plates and read directly for NLuc activity (Promega) on a Synergy HT plate reader (BioTek). Multicycle experiments were seeded in clear bottom plates and observed for siRNA toxicity. Supernatants were collected and titrated by infecting MDCK cells (white bottom 96-well plates) for 1 h with 20 µl supernatant, washing twice with VGM, and incubating for 8 h. Luciferase activity was read as above. Single cycle infections were also performed with CA04 (MOI, 0.5) and B/Bris (MOI, 1) PASTN viruses as above. Virus gene expression (single cycle infection) and titer (multicycle infection) were normalized to non-targeting siRNA control. For knockdown during wildtype virus infection, A549 cells were forward transfected for two days with 25 nM siRNA in 24-well plates and infected with WSN (MOI, 0.01) for 24 h. Supernatants were titrated by plaque assay on MDCK cells.

For overexpression experiments, 293T cells were reverse transfected with hnRNP UL1, NXF1, or TPR using TransIT-2020 (Mirus) in 96-well plates for 24 h. Transfected cells were infected with PASTN virus at an MOI of 0.01 for 24 h, and supernatants titrated as above. A549 cells stably overexpressing hnRNP UL1, MECR or cMECR, or MECR knockout cells overexpressing Etr1-HA were clonally isolated and analyzed for V5 or HA-tagged protein expression, respectively. Etr1-complemented cells were then transduced with cMECR-containing lentiviruses and polyclonal blasticidin-resistant cells used for experimental analysis. A549 overexpression cells were infected with PASTN virus in 96-well plates for 24 h at an MOI of 0.05. Supernatants were titrated as above.

Infection experiments analyzing the role of mtFAS were performed by treating A549 cells with siRNAs for 48 h in 96-well plates with 50 nM non-targeting control siRNA, 25 nM non-targeting control mixed with 25 nM MECR siRNA, 25 nM non-targeting control mixed with 25 nM *NDUFAB1* (ACP) siRNA, or 25 nM of MECR and ACP siRNAs, followed by multicycle analysis with WSN PASTN as described above. For drug treatment, A549 cells were seeded in 96-well plates and treated with DMSO or 50 µM C75 (Sigma C5940) for 24 h, infected with WSN PASTN (MOI, 0.05; 24 h) in the presence of DMSO or 50 µM C75, and titrated as above.

### Co-immunoprecipitations

Interactions between endogenous hnRNP UL1 and PB2 were tested by co-immunoprecipitation. A549 cells were infected with WSN PB2-FLAG (10 cm dish; MOI, 1; 18 h), lysed in co-IP buffer, lysates immunoprecipitated overnight with anti-FLAG resin (M2, Sigma), and co-precipitating hnRNP UL1 was detected by western blot. For V5 immunoprecipitations, mammalian expression plasmids encoding PB2-FLAG, PB1-FLAG, PA, and HNRNPUL1-V5 or MECR-V5 were forward transfected into 293T cells using PEI (PEI MAX, Polysciences; 6-well plates). Cells were lysed in co-IP buffer, immunoprecipitated with anti-V5 antibody, and co-precipitating viral proteins were detected by western blot. To test the effect of viral RNA on polymerase:hnRNP UL1 interactions, plasmids expressing vNA or NP were included where indicated. Cells were lysed two days post-transfection in co-IP buffer with or without 1 µl RNase A (Thermo Scientific). Interactions with WT MECR during infection were tested by forward transfecting 293T cells with mammalian expression plasmids encoding PB2-FLAG, PB1-FLAG, PA, and MECR-V5 or cMECR-V5 for 42 h and infecting with WSN (MOI 10; 6 h) followed by lysis. Clarified lysates were incubated with 1.5 µg polyclonal anti-V5 or IgG for 1 h and captured with protein A agarose resin (P2545, Sigma) for 30 min. Immunoprecipitates were recovered, washed four times with co-IP buffer and eluted by boiling in Laemmli sample buffer. Interactions with codon-optimized MECR- and cMECR-mCh fusions were tested by transfecting 293T cells with mammalian expression constructs using TransIT-X2 (Mirus) for 48 h. Where indicated, plasmids expressing PB2-FLAG, PB1-FLAG, PA, NP and genomic RNA vNA-GFP were co-transfected. Cells were lysed in co-IP buffer, mCherry fusion proteins were captured with RFP-Trap (Chromotek RTA-20) for 3 h at 4° C, and resin was washed three times with co-IP buffer prior to elution by boiling in Laemmli sample buffer. Samples were separated by SDS-PAGE and analyzed by western blotting. Chemiluminescent images were captured on an Odyssey Fc Imager and quantified using Image Studio v5.2.5 (LI-COR). At least two biological replicate experiments were performed.

For concurrent analysis of cMECR interactions and RNP assembly, TransIT-X2 was used to transfect 293T cells with plasmids expressing codon-optimized MECR-mCh, cMECR-mCh or mCherry alone, along with PB2-FLAG, PB1-FLAG, PA, and the genomic RNA vNA-GFP in the presence or absence of NP-V5. Cells were lysed with co-IP buffer 2 d post-transfection and divided in half to assess interactions with cMECR using RFP-Trap or RNP assembly using V5-Trap (Chromotek V5TA-20). Samples were rocked for 3 h at 4 ° C. Immunoprecipitates were recovered, washed once with 500 mM NaCl co-IP buffer, then twice more with co-IP buffer, and finally eluted by boiling in Laemmli sample buffer. Samples were separated by SDS-PAGE and analyzed by western blotting.

Proximity ligation utilized the biotin ligase AirID^53^. Assays were performed by expressing polymerase proteins in 293T cells with free AirID or AirID fused to the C-terminus of codon-optimized cMECR-V5. Exogenous biotin [50 µM] (Sigma B4501) was added 18 h posttransfection. Lysates were prepared at 48 h and PB2 was immuno-precipitated, blotted and probed with streptavidin-HRP (LI-COR 925-32230) or PB1 antibodies. Input samples were additionally probed with PB2 and V5 antibodies.

### Fluorescence microscopy

For immunofluorescence, 293T cells were transfected in 48-well plates with plasmids expressing V5-tagged hnRNP UL1, MECR or cMECR and 48 h later infected with WSN PB2-FLAG (MOI, 3; 8 h). A549 cells stably expressing hnRNP UL1 or MECR were seeded on coverslips in 12-well plates and infected with WSN PB2-FLAG (MOI, 0.5; 8 h). Monolayers were fixed with 4% paraformaldehyde in PBS for 10 min, quenched and permeabilized with 0.1 M glycine + 0.1% Triton X-100 in PBS for 5 min, and blocked with 3% BSA in PBS for 30 min at room temperature. Primary and secondary antibodies were sequentially incubated for 1 h each at room temperature at 1 µg/ml in blocking buffer. DAPI was added to 293T cells during secondary antibody incubation. Coverslips were mounted in medium containing DAPI (Vector Laboratories, H-1200). Images were captured using a 20X objective on an EVOS FL Auto (ThermoFisher) and processed in Adobe Photoshop CC.

For live cell imaging, 10X or 20X images were captured on an EVOS FL Auto one day post transfection. To visualize nuclei, Hoechst 33342 (Anaspec AS-83218) was added 10 minutes prior to capture. GFP-positive cells were enumerated using Image J by automating the following commands: threshold (300); binary > watershed; analyze particles (size 100-500 px).

### RNA-sequencing

RNA was isolated from A549 cells that were mock treated, interferon-β treated (250 U/ml for 8 h) or infected with influenza WSN (MOI 0.02 for 24 h) using TRIzol (Invitrogen). Biologic triplicate sample RNA was sent for library preparation and paired-end RNA-seq by Novogene (BioProject PRJNA667475). Sequences were trimmed with BBDuk in the BBMap suite and aligned to the human genome (hg38) with HISAT2^88,89^. Splicing events in the MECR locus were visualized in IGV (sashimi plot function) to enumerate the exon-joining reads into the 5’ boundary of exon 2^90^.

### cMECR initiation site identification and splicing reporter

Ribosome profiling data were acquired from Lee et al.^51^ In that study, HEK293 cells were treated with 50 µM lactimidomycin (LTM) for 37 °C for 30 min to arrest ribosomes at start sites prior to the first elongation event. We trimmed ribosome protected fragments from Illumina HiSeq 2000 runs (SRR618772, LTM rep1; SRR618773, LTM rep2) with BBDuk in the BBMap suite and aligned to the human genome (hg38) with HISAT2^88,89^. Reads mapping to the *MECR* locus exon 1-2 were visualized with IGV.

cMECR splice reporters were constructed by inserting portions of the *MECR* locus upstream of the cMECR-V5 ORF. The native intron is ∼14,000 bp, therefore a miniaturized version was used for the “Splice” reporter. 80 nt from the 3’ of exon 1 and 100 nt from the 5’ end of intron 1 were fused 100 nt upstream of the cMECR splice acceptor site, leading into the remainder of the cMECR UTR and the cMECR start codon (M77 in MECR). The “Splice+FLAG” reporter was then created by replacing the fourth uORF with a start site and coding sequence for a FLAG epitope that remained in frame with cMECR-V5.

### Phylogenetic analysis

MECR amino acid sequences were retrieved from NCBI (**Supp. Table 4**). Phylogenetic analysis was performed using Influenza Research Database (www.fludb.org) using PhyML options^91^ to generate a Newick file. FigTree v1.4.4 (http://tree.bio.ed.ac.uk/software/figtree) was used for visualization.

### Statistical Analysis

Assays were performed with three to six technical replicates and represent at least three independent biological replicates. Mean and SD or SEM were calculated. When data were normalized, error was propagated to each individual experimental condition. Statistical significance was determined for pairwise comparisons with a Student’s *t* test and for multiple comparisons an ANOVA and *post hoc* Dunnett’s or Sidak’s test. Correlation across data sets was determined via a two-tailed Pearson correlation coefficient. Values from statistical analyses of siRNA screens are reported (**Supp Table 3**). Statistical tests were performed using Prism (v 8.4.2, GraphPad).

