## Supplementary material for "Alternative splicing liberates a cryptic cytoplasmic isoform of mitochondrial MECR that antagonizes influenza virus": Supp. Fig. 1

**Supp. Fig. 1: Validation of ICC-MS and network analysis. a**, Relative protein abundance of PB2 (red), PA (cyan), and NP (yellow) in PB2 ICC-MS samples shows decreasing capture with increasing competition antibody. Data shown are in biological triplicate. **b**, Fully annotated and zoomable version of the diagram in Fig 1d. Minimum-cost flow simulations connect top PB2 interactors identified by ICC-MS (red) to influenza host factors (gray) through novel host proteins (white). Modules comprising different PB2 interactors with enriched GO terms and *P* values indicated. Node sizes indicate empirical *P* values based on the control flow simulation. **c**, Enlarged view of MECR-containing module. **d**, Validation shows that flow simulation networks of protein-protein interactions reflect biochemical pathways that regulate viral replication. Flow simulation captured PKR (*EIF2AK2*) interactions with RNA binding proteins DHX30, IGF2BP2, and TARBP, and linked RIG-I (*DDX58*) with TRIM14 through MAVS, and also IFIT3, LGP2, and USP15. Additionally, mRNA nuclear export pathways were faithfully reconstructed with NXF1 connecting with NXT2 and CRM1 (*XPO1)* connecting with NXF3. Our networks linked EXOSC3 to the exosome components EXOSC4, EXOSC5, and EXOSC8 and with the NEXT accessory complex (*RBM7*), both of which are important for influenza transcription [4].

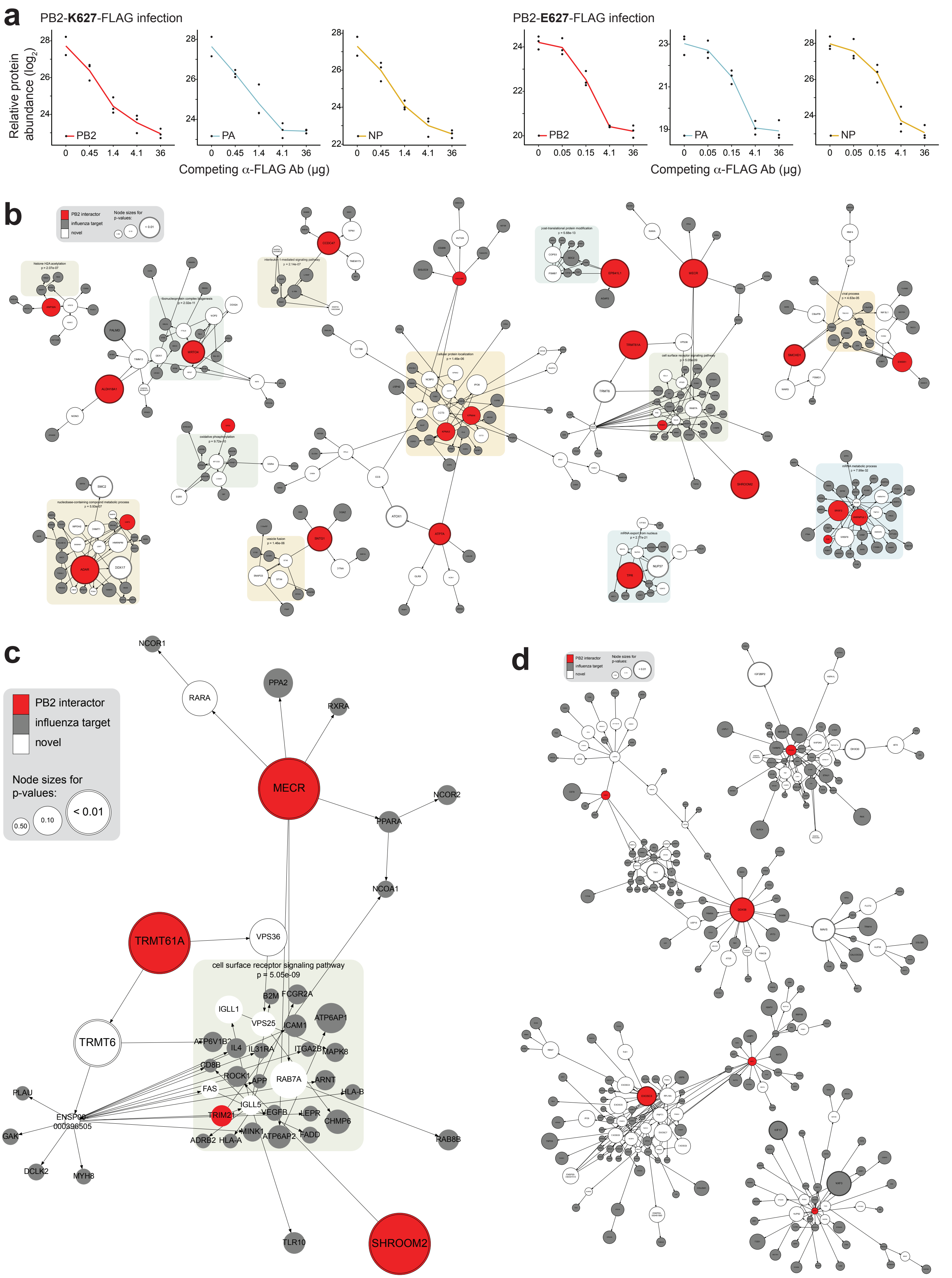
a
PB2-K627-FLAG infection
PB2-E627-FLAG infection
Relative protein abundance (log2)
PB2
PA
NP
Competing α-FLAG Ab (μg)
0
0.45
1.4
4.1
36
0.05
0.15
b
PB2 interactor
influenza target
novel
Node sizes for p-values:
< 0.01
c
PB2 interactor
influenza target
novel
Node sizes for p-values:
0.50
0.10
< 0.01
NCOR1
RARA
PPA2
RXRA
MECR
NCOR2
PPARA
NCOA1
TRMT61A
VPS36
cell surface receptor signaling pathway
p = 5.05e-09
B2M
FCGR2A
IGLL1
VPS25
ICAM1
ATP6AP1
TRMT6
ATP6V1B2
IL4
IL31RA
ITGA2B
MAPK8
CD8B
ROCK1
APP
RAB7A
ARNT
FAS
HLA-B
PLAU
IGLL5
VEGFB
LEPR
ENSP00000396505
TRIM21
CHMP6
GAK
MINK1
FADD
ADRB2
HLA-A
ATP6AP2
RAB8B
DCLK2
MYH8
TLR10
SHROOM2
d
PB2 interactor
influenza target
novel
Node sizes for p-values:
< 0.01
