## Supplementary material for "Alternative splicing liberates a cryptic cytoplasmic isoform of mitochondrial MECR that antagonizes influenza virus": Supp. Fig. 2

**Supp. Fig. 2: Functional analyses of top candidate PB2 interactors.** **a**, Secondary screening of proteomic hits by siRNA treatment and reporter virus infection. After knockdown, 293T cells were infected with human (PB2-627K; MOI, 0.01) or avian-adapted (PB2-627E; MOI, 0.05) WSN NLuc virus for 24 h. Viral supernatants were titered and normalized to a non-targeting control (NT). Control NXF1 (gray) and validated proteins highlighted (hnRNP UL1, cyan; MECR, yellow). Data are mean ± SEM of *n* = 2-3 biological replicates. **b**, Concordance of virus titer for PB2-627E *vs* PB2-627K virus infections in siRNA-treated cells (from **a**). Statistical analysis performed with a two-tailed Pearson correlation coefficient. **c**, Screening as in **a**, except A549 cells were infected with viruses as above for a single-cycle of infection (MOI, 0.1; 8 h) and virus gene expression in infected cells normalized as above. **d**, Concordance of gene expression (data from **c**, analyzed as in **b**).

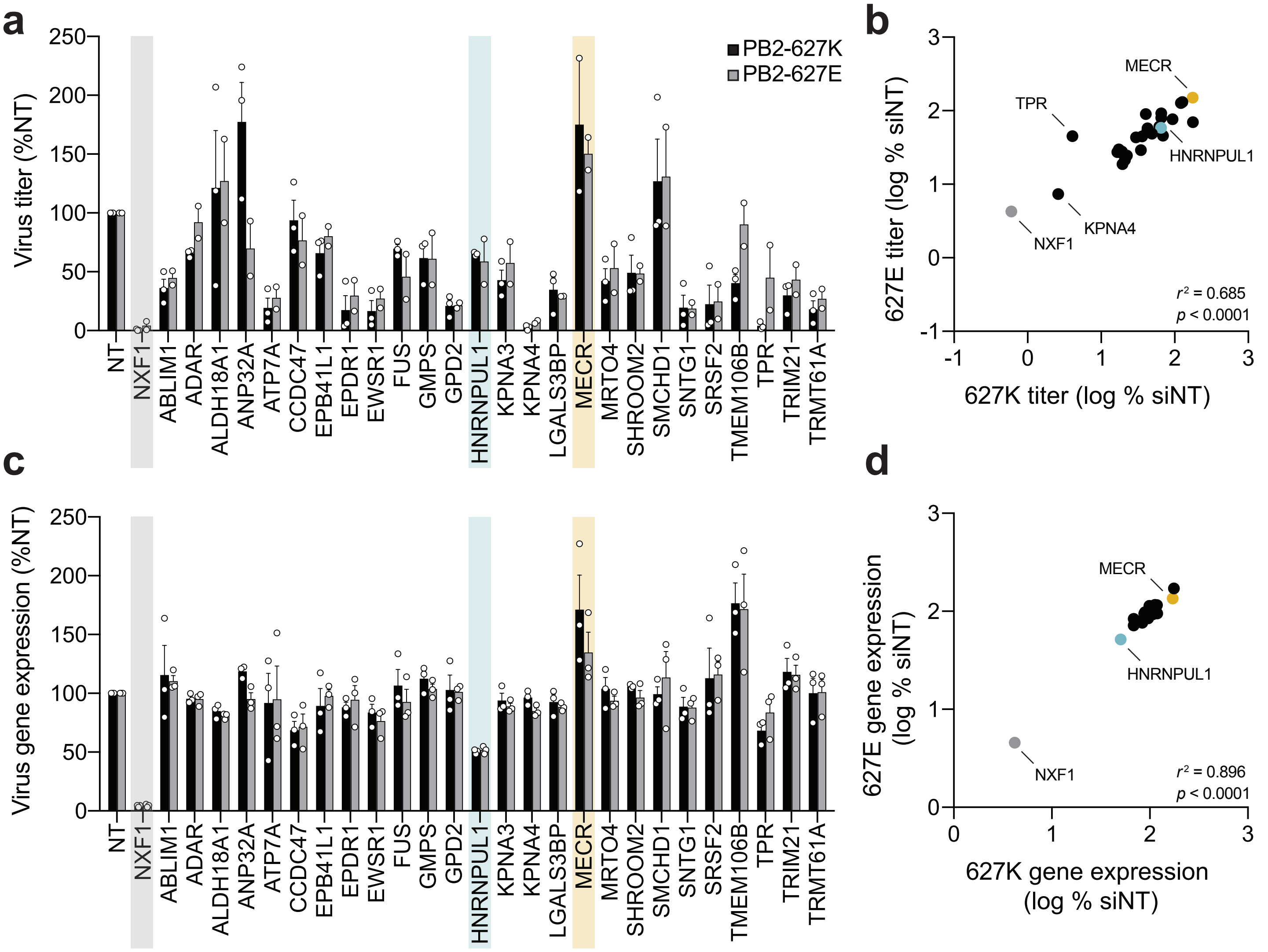
a
b
c
d
PB2-627K
PB2-627E
Virus titer (%NT)
250
200
150
100
50
0
NT
NXF1
ABLIM1
ADAR
ALDH18A1
ANP32A
ATP7A
CCDC47
EPB41L1
EPDR1
EWSR1
FUS
GMPS
GPD2
HNRNPUL1
KPNA3
KPNA4
LGALS3BP
MECR
MRTO4
SHROOM2
SMCHD1
SNTG1
SRSF2
TMEM106B
TPR
TRIM21
TRMT61A
627E titer (log % siNT)
627K titer (log % siNT)
MECR
TPR
HNRNPUL1
KPNA4
NXF1
$r^2$ = 0.685
$p$ < 0.0001
-1
0
1
2
3
Virus gene expression (%NT)
627E gene expression (log % siNT)
627K gene expression (log % siNT)
$r^2$ = 0.896
$p$ < 0.0001
