## Supplementary material for "Alternative splicing liberates a cryptic cytoplasmic isoform of mitochondrial MECR that antagonizes influenza virus": Supp. Fig. 3

**Supp. Fig. 3: Enhanced expression of nuclear mRNA export proteins increases virus production.** **a**, Replication of WSN NLuc (MOI, 0.05; 24 h) was measured in 293T cells transfected with empty vector (EV) or with plasmids encoding the indicated RNA export proteins. Mean ± SD of $n = 3$. One-way ANOVA with *post hoc* Dunnett's multiple comparisons test; ****, $P < 0.0001$. **b**, Immunofluorescence microscopy of WT and mutant hnRNP UL1 expressed in 293T cells, scale bars, 20 μm.

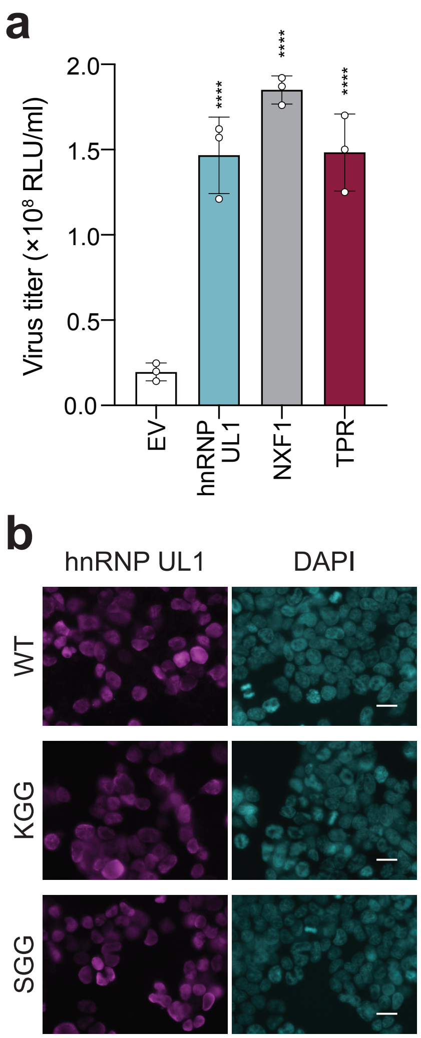

a
Virus titer (×10^8 RLU/ml)
2.0
1.5
1.0
0.5
0.0
****
****
****
EV
hnRNP UL1
NXF1
TPR
b
hnRNP UL1
DAPI
WT
KGG
SGG
