## Supplementary material for "Alternative splicing liberates a cryptic cytoplasmic isoform of mitochondrial MECR that antagonizes influenza virus": Supp. Fig. 4

**Supp. Fig. 4: Allelic identities of *MECR* knockout A549 cells.** Deep sequencing of clonal knockout cells. CRISPR gRNA sequence underlined, PAM dotted. In-frame amino acids indicated on top. KO-1 is homozygous and KO-2 has two distinct alleles.

```
          V  G  G  N  E  G  V  A  Q  V  V  A   V  G  S  N  V  T  G
WT        GTTGGAGGGAACGAAGGTGTTGCACAGGTGGTAGC-GGTGGGCAGCAATGTGACCGGG
KO-1      GTTGGAGGGAACGAAGGTGTTGCACAGGTGGTAGCAGGTGGGCAGCAATGTGACCGGG
KO-2a     GTTGGAGGGAACGAAGGTGTTGCACAGGTGGTAGCGGGTGGGCAGCAATGTGACCGGG
KO-2b     GTTGGAGGGAACGAAGGTGTTGCACAGGT-----------GGTAGCAATGTGACCGGG
                                             ^
```
