## Supplementary material for "Alternative splicing liberates a cryptic cytoplasmic isoform of mitochondrial MECR that antagonizes influenza virus": Supp. Fig. 5

**Supp. Fig. 5: Characterization of endogenous *MECR* transcripts and cMECR:polymerase interactions. a-b**, RT-PCR analysis of poly-adenylated *MECR* transcripts in mock or WSN-infected A549 cells (MOI 1, 6 h). **a**, Genomic representation of *MECR* and regions amplified by specific primer sets. Untranslated regions (UTRs) in dark gray boxes, exons in light gray boxes, and introns in lines. Diagram segments are not to scale; base pair (bp) lengths are indicated. **b**, Amplification of transcripts containing exon 1 (red), a region spanning exon 1 and 2 (green), and the unique cMECR UTR through exon 3 (yellow). The primer pair MECR ex1 FOR/MECR ex2 REV amplifies MECR (205 bp, empty circle) and cMECR (332 bp, filled circle). cMECR is predicted to be variable lengths around 332 bp due to the use of multiple splice donor sites paired with the cMECR UTR splice acceptor site. Asterisk; non-specific band. Size shown in bp derived from a DNA ladder. **c**, Ribosomes initiate on the short splice isoform transcript in the UTR and at the cMECR start site. Cells were treated with lactimidomycin to identify initiation sites during ribosome profiling (data from [51]). Ribosome protected fragments (RPF) and total RNA (RNA-Seq) were mapped to the MECR locus, showing initiation events at the predicted cMECR start codon and other sites in the short splice isoform encoding cMECR. Zoom on exon 2 shows coding potential of the short isoform with stop codons in red, start codons in green, and the cMECR initiation site with an arrow. **d,** PB2-FLAG and V5-tagged MECR or cMECR were expressed in 293T cells with or without the other polymerase (PB1/PA) or vRNP (vNA/NP) components. Cells were lysed and immunoprecipitated with anti-V5 antibody or IgG controls. Proteins were detected by western blot. **e**, Subcellular localization of mCherry (mCh) fusion constructs. Plasmids encoding codon-optimized MECR (M77L) or cMECR fused to mCh with a 5GS linker were transfected into 293T cells. Red fluorescence images were captured 24 h post transfection. Scale bars; 20 µm. **f**, cMECR-AirID biotinylates polymerase. Cells co-expressing PB2-FLAG tagged polymerase and free AirID or the cMECR-AirID fusion were immuno-precipitated with anti-FLAG. Input and co-precipitating proteins were detected by western blotting with streptavidin-HRP or antibodies recognizing the indicated proteins.

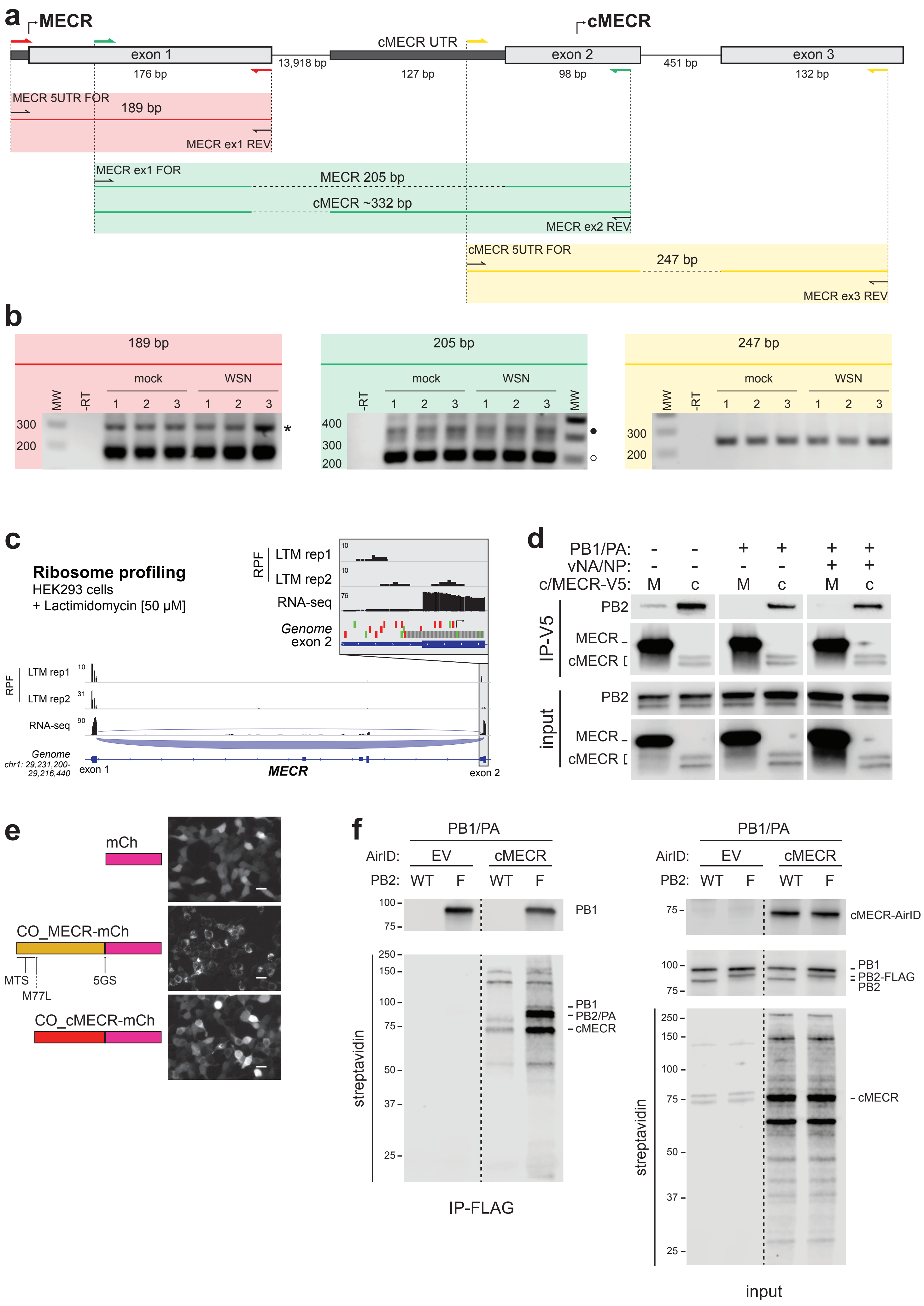
a
MECR
cMECR
exon 1
176 bp
13,918 bp
cMECR UTR
127 bp
exon 2
98 bp
451 bp
exon 3
132 bp
MECR 5UTR FOR
189 bp
MECR ex1 REV
MECR ex1 FOR
MECR 205 bp
cMECR ~332 bp
MECR ex2 REV
cMECR 5UTR FOR
247 bp
MECR ex3 REV
b
189 bp
205 bp
247 bp
mock
WSN
MW
-RT
1 2 3 1 2 3
300
200
400
c
Ribosome profiling
HEK293 cells
+ Lactimidomycin [50 μM]
RPF
LTM rep1
LTM rep2
RNA-seq
Genome
exon 2
exon 1
MECR
chr1: 29,231,200-29,216,440
d
PB1/PA:
vNA/NP:
c/MECR-V5:
M c M c M c
IP-V5
input
PB2
MECR
cMECR
e
mCh
CO_MECR-mCh
MTS
M77L
5GS
CO_cMECR-mCh
f
PB1/PA
AirID:
EV
cMECR
PB2:
WT F WT F
PB1
PB2/PA
cMECR
cMECR-AirID
PB2-FLAG
PB2
streptavidin
IP-FLAG
input
250
150
100
75
50
37
25
