## Supplementary material for "Alternative splicing liberates a cryptic cytoplasmic isoform of mitochondrial MECR that antagonizes influenza virus": Supp. Fig. 6

**Supp. Fig. 6: cMECR binds to and shuts down polymerase activity**. **a**, Representative images used for GFP-positive cell counting in Fig. 6d. GFP-based polymerase activity assays were performed in 293T cells co-expressing mCh, MECR M77L-mCh, or cMECR-mCh. NP was omitted in the negative control (-NP). *Top*: 10X whole-field view of vNA-GFP fluorescence. *Bottom*: Zoomed view with underlying Hoechst and mCh fluorescence. Dotted box indicates zoomed section. Scale bars for both images are 200 μm. **b**, Etr1-expressing *MECR* knockout cells were transduced with lentivirus expressing cMECR or an empty vector control. Cells were subsequently infected with WSN NLuc (MOI, 0.05; 24 h) and viral titers were measured in supernatants. Mean ± SD of $n = 3$. Two-way ANOVA with *post hoc* Dunnett's multiple comparisons test to compare to untransduced cells; *, $P < 0.05$; **, $P < 0.01$. **c**, MECR antiviral effect does not require innate immune signaling pathways. siRNA-treated WT or *PKR, RIG-I*, or *MAVS* knockout A549 cells were infected with WSN NLuc (MOI, 0.05). Viral titers were measured 24 h post infection and normalized to NT controls in the corresponding knockout line. Mean ± SD of $n = 3$. Two-way ANOVA with *post hoc* Dunnett's multiple comparisons test to compare to WT cells; ****, $P < 0.0001$; other conditions were not significant.

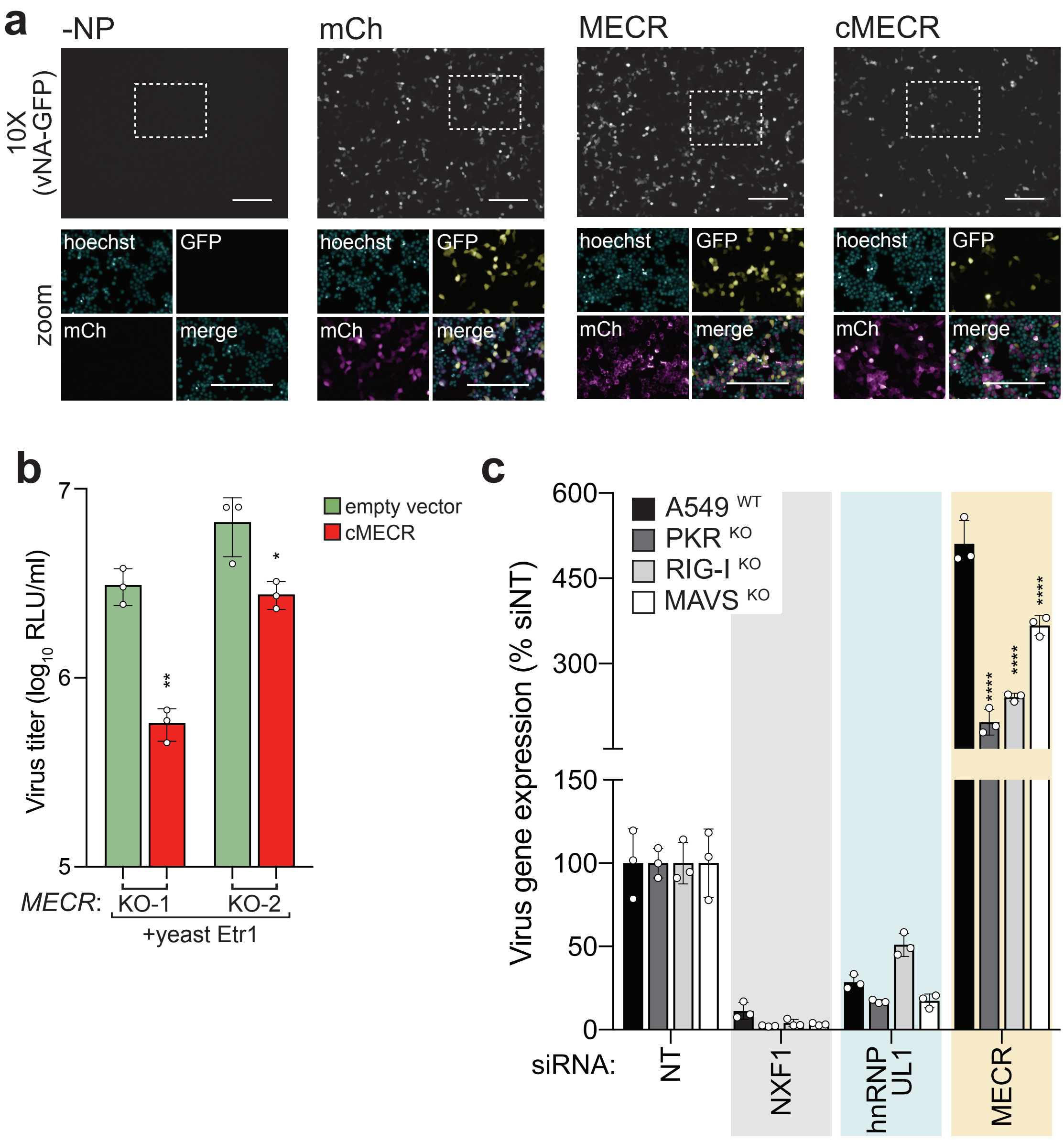

a
-NP
mCh
MECR
cMECR
10X (vNA-GFP)
zoom
hoechst
GFP
mCh
merge
b
Virus titer (log10 RLU/ml)
7
6
5
empty vector
cMECR
**
*
MECR:
KO-1
KO-2
+yeast Etr1
c
Virus gene expression (% siNT)
600
450
300
150
100
50
0
A549 WT
PKR KO
RIG-I KO
MAVS KO
****
****
****
siRNA:
NT
NXF1
hnRNP UL1
MECR
