## Supplementary material for "Alternative splicing liberates a cryptic cytoplasmic isoform of mitochondrial MECR that antagonizes influenza virus": Supp. Table 4

**Supp. Table 4.** Accession numbers for MECR homologs

| Common Name | Species | MECR[a] | cMECR[a] |
|---|---|---|---|
| Human | Homo sapiens | NP_057095.4 | NP_001336643.1 |
| Chimpanzee | Pan paniscus | XP_003828155.1 | XP_034815583.1 |
| Gorilla | Gorilla gorilla gorilla | XP_004025368.3 | XP_018869404.1 |
| Rhesus macaque | Macaca mulatta | NP_001248098.1 | XP_014988731.1 |
| Pale spear-nosed bat | Phyllostomus discolor | XP_028368816.1 | |
| American bison | Bison bison bison | XP_010828514.1 | |
| Cow | Bos taurus | NP_858055.1 | XP_024855139.1 |
| Pig | Sus scrofa | NP_001231011.1 | XP_020949000.1 |
| Horse | Equus caballus | XP_001503984.3 | |
| Beluga whale | Delphinapterus leucas | XP_022407680.2 | |
| Camel | Camelus ferus | XP_006175527.1 | XP_032351592.1 |
| Alpaca | Vicugna pacos | XP_031539466.1 | |
| Mouse | Mus musculus | NP_079573.2 | XP_036020031.1 |
| Chicken | Gallus gallus | XP_024998883.1 | |
| Goose | Anser cygnoides domesticus | XP_013050980.1 | XP_013050980.1 |
| Duck | Anas platyrhynchos | XP_027299737.1 | |
| Saker falcon | Falco cherrug | XP_027668811.1 | XP_005433096.1 |
| Golden eagle | Aquila chrysaetos chrysaetos | XP_029895600.1 | XP_029895602.1 |
| Alligator | Alligator mississippiensis | KYO48103.1 | XP_019344297.1 |
| Salmon | Salmo salar | XP_014055905.1 | |
| Trout | Salmo trutta | XP_029592657.1 | |
| Zebrafish | Danio rerio | AAI53449.1 | |
| Frog | Xenopus tropicalis | NP_001016371.1 | XP_012812419.1 |
| Slime mold | Dictyostelium fasciculatum | XP_004362847.1 | |
| Nematode | Trichinella spiralis | XP_003380885.1 | |
| Baker's yeast | Saccharomyces cerevisiae | NP_009582.1 | |

*a*, NCBI Reference Sequence ID
