## Supplementary material for "Alternative splicing liberates a cryptic cytoplasmic isoform of mitochondrial MECR that antagonizes influenza virus": Supp. Table 3

**Supp. Table 3.** Statistical analysis[a] of influenza infection siRNA screens

| | Fig. 2a-b[b] | | Supp. Fig. 2a-b[c] | | Supp Fig. 2c-d[d] | |
|---|---|---|---|---|---|---|
| siRNA target | *P* value | sig[e] | *P* value | sig[e] | *P* value | sig[e] |
| NXF1 | 0.003 | ** | <0.0001 | **** | <0.0001 | **** |
| ABLIM1 | 0.0932 | | 0.0013 | ** | 0.2818 | |
| ADAR | 0.7809 | | 0.2433 | | 0.6918 | |
| ALDH18A1 | 0.2871 | | 0.1801 | | 0.1555 | |
| ANP32A | 0.4756 | | 0.1942 | | 0.5698 | |
| ATP7A | 0.7944 | | <0.0001 | **** | 0.5782 | |
| CCDC47 | 0.4571 | | 0.4107 | | 0.014 | * |
| EPB41L1 | 0.251 | | 0.1353 | | 0.5914 | |
| EPDR1 | 0.1315 | | <0.0001 | **** | 0.4658 | |
| EWSR1 | 0.2701 | | <0.0001 | **** | 0.0986 | |
| FUS | 0.4095 | | 0.0205 | * | 0.9769 | |
| GMPS | 0.9369 | | 0.0337 | * | 0.5055 | |
| GPD2 | 0.1871 | | <0.0001 | **** | 0.8706 | |
| HNRNPUL1 | 0.0319 | * | 0.0363 | * | <0.0001 | **** |
| KPNA3 | 0.3599 | | 0.0067 | ** | 0.4731 | |
| KPNA4 | 0.0906 | | <0.0001 | **** | 0.4336 | |
| LGALS3BP | 0.7595 | | 0.0003 | *** | 0.4312 | |
| MECR | <0.0001 | **** | 0.0013 | ** | <0.0001 | **** |
| MRTO4 | 0.9697 | | 0.0045 | ** | 0.9172 | |
| SHROOM2 | 0.7592 | | 0.0054 | ** | 0.93 | |
| SMCHD1 | 0.7206 | | 0.1106 | | 0.5983 | |
| SNTG1 | 0.5135 | | <0.0001 | **** | 0.3284 | |
| SRSF2 | 0.459 | | <0.0001 | **** | 0.2338 | |
| TMEM106B | 0.7947 | | 0.0562 | | <0.0001 | **** |
| TPR | 0.3615 | | <0.0001 | **** | 0.047 | * |
| TRIM21 | 0.4499 | | 0.0007 | *** | 0.1607 | |

*a*, Two-way ANOVA followed by Fisher's LSD test
*b*, A549 multicycle
*c*, 293T multicycle
*d*, A549 single cycle
*e*, ****, $P < 0.0001$; ***, $P < 0.001$; **, $P < 0.01$; *, $P < 0.05$
